# OptoAssay – Light-controlled Dynamic Bioassay Using Optogenetic Switches

**DOI:** 10.1101/2021.11.06.467572

**Authors:** Nadine Urban, Maximillian Hörner, Wilfried Weber, Can Dincer

## Abstract

Circumventing the limitations of current bioassays, we introduce the first light-controlled assay, the OptoAssay, towards wash- and pump-free point-of-care diagnostics. Extending the capabilities of standard bioassays with light-dependent and reversible interaction of optogenetic switches, OptoAssays enable a bi-directional movement of assay components, only by changing the wavelength of light. Combined with smartphones, OptoAssays obviate the need for external flow control systems like pumps or valves and signal readout devices.

## Main

Point-of care (POC) diagnostics is the key player for fast interventions that might have critical value for patient’s outcome. For this purpose, POC testing that allows for rapid diagnostics in non-laboratory settings carried out by untrained personnel has become more commonplace over the last decade. Its importance has even more deepened during the ongoing Covid19 pandemic. One of the most used POC formats are paper-based devices like lateral flow assays (LFAs), where the sample is added on a cellulose-based test stripe and transported through capillary forces along the stripe. This, however, only allows for a uni-directional sample flow which fundamentally limits the flexibility in assay design and sample processing^1,2^. On the other hand, other POC devices offering bidirectional microfluidics suffer from expensive and bulky pumps or flow control systems, also requiring an additional energy source, that complicates their on-site use^3,4^.

To circumvent this restraint, we present for the first time the proof-of-concept of a light-controlled dynamic bioassay (OptoAssay). OptoAssay allows for bi-directional movement of biomolecules, enabling wash-free signal readout of bioassays. For this purpose, the biomolecules fused to a phytochrome interacting factor 6 (PIF6) can be released from and rebind to the plant photoreceptor phytochrome B (PhyB) by exposure to far-red or red light, respectively, thereby circumventing the need for external flow control devices. PhyB exhibits two distinct light absorbing conformations (**Figure 1A**): a red-light absorbing P_r_ (red) state with λ_max_ ~ 651 nm^5^ nm and a far-red light absorbing, biologically active, P_fr_ (far red) state with λ_max_ ~ 713 nm. In the active state, PhyB can bind proteins from a class called phytochrome interacting factors while in the inactive state this interaction is reversed^6,7^. Up to date, this switching behavior of the PhyB/PIF systems has been used in various applications to control, for example, gene expression^8^, cell signalling^9^ or to create biohybrid materials^10^.

**Figure 1.**
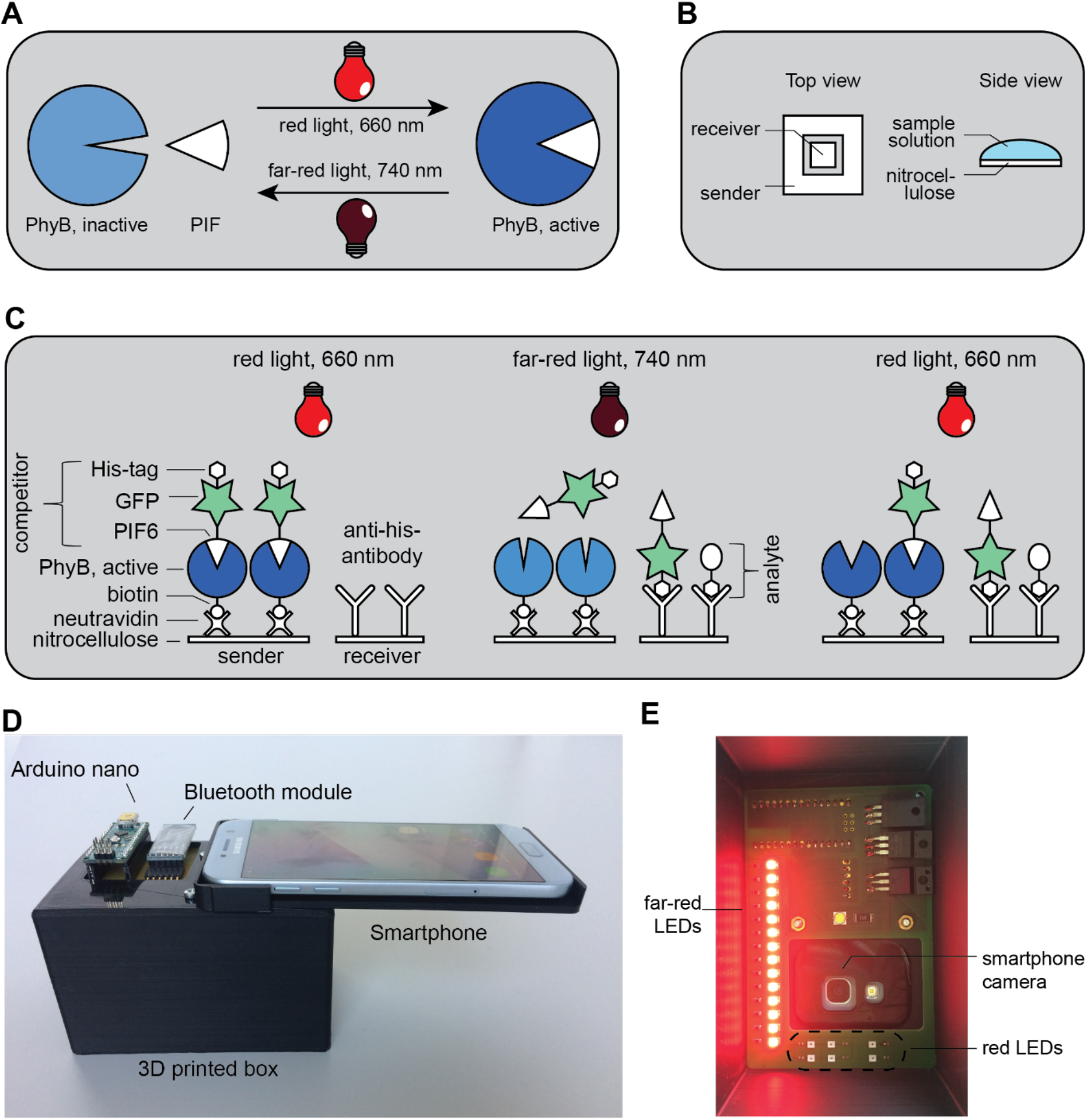
**A)** Scheme describing the basics of Phytochrome B (PhyB) and Phytochrome Interacting Factor (PIF) interaction. PhyB is transformed in the active form by illumination with red light (660 nm) where it interacts with PIF. By exposure to far-red light (740 nm), PhyB is converted to the inactive form where it cannot interact with PIF. **B)** Schematics describing the design of the OptoAssay. The smaller square-shaped receiver area is enclosed by a larger square-shaped sender area. Both zones are connected with sample solution on top of the membranes. **C)** Working principle of the OptoAssay. In the initial state, active PhyB is immobilized via neutravidin (nAv) on the nitrocellulose sender area while the competitor complex (PIF6-GFP-his) is associated with PhyB during red light illumination. The receiver area is coated with anti his-tag antibodies. In the second step, where the assay is illuminated with far-red light and the competitor is released from the inactive PhyB conformation while at the same time an analyte molecule (his-tagged protein) is added. Analyte and competitor compete now for binding to the antibodies on the receiver area. The final step comprises of illumination with red light in order to reassociate all unbound competitors to PhyB on the sender area. **D)** 3D printed PhotoBox that allows for sample illumination and signal readout for the envisioned POC scenario. A smartphone is placed on top of the box and connected via a Bluetooth module to an Arduino nano to switch on and off the illumination. Through a hole in the box, images can be taken with the smartphone camera. **E)** PhotoBox from inside: LEDs for far-red- and red-light illumination.

Here, we apply optogenetic switches for realizing the first light-controlled dynamic bioassay on a nitrocellulose substrate and successfully demonstrate its proof-of-principle, detecting a hexahistidine-tag (referred to as his-tag) as analyte. Besides, we show the general versatility of the assay components on other substrates, namely agarose beads and poly(methyl methacrylate) (PMMA), employed for building POC devices.

## Results

The concept of the OptoAssay consists of two distinct areas (**Figure 1B**): a smaller receiver area where the signal readout is performed, is placed in the middle of the sender area into a notch so that the areas are not directly connected. These areas are later bridged by adding sample solution to both sides so that the assay components can diffuse from one area to the other. In this work, the OptoAssay is carried out as an immunoassay using antibodies. Herein, a competitive assay format was chosen, in which a competitor and the analyte in the sample compete for binding to the detection antibody. The sender area (**Figure 1C**) is treated with neutravidin (nAv) so that the light sensitive components PhyB can be immobilized via a biotin-neutravidin interaction. Here, PIF6 can bind to PhyB or be released depending on the wavelength applied. On the other hand, antibodies directed against the molecule of interest are immobilized on the receiver area.

For the proof-of-principle assay, we use a his-tagged protein (see **Methods**) as analyte and his-tagged PIF6 as competitor, both of which are recognized by the anti his-tag antibody immobilized in the receiver area. The signal readout is performed by measuring the fluorescence intensity of the competitor complex, his-tagged PIF6, to which green fluorescent protein (GFP) is fused. For the initial OptoAssay configuration, the competitor complex is attached to PhyB on the sender area during red light exposure. The assay procedure itself comprises of two steps (**Figure 1B**): first, far-red light illumination is used to release the competitor complex from PhyB. At the same time, the analyte is added. Now both the analyte and the competitor can compete for binding to the antibody immobilized on the receiver area. Second, red light is applied to re-associate unbound competitor complexes in the sender area and thus, to remove them from the receiver area. Due to the competitive assay format, the signal measured on the receiver area is inverse proportional to the concentration of the analyte. In order to enable POC testing using the OptoAssay, a 3D printed PhotoBox that allows for illumination with red and far-red light has been built (**Figure1 D,E**)^11^. The illumination can be controlled with a smartphone that is linked via a Bluetooth module to the electronics of the PhotoBox. Images of the OptoAssay results can be then easily read out, evaluated, and further transmitted to, for example, medical facilities using a smartphone.

As substrate material for immobilization of assay components, we chose nitrocellulose since it is widely applied for LFA devices due to its non-specific affinity to proteins^12,13^. We first investigated an appropriate blocking method for nitrocellulose to cover unoccupied spaces after the protein of interest has been immobilized. Therefore, we examined the blocking performance of bovine serum albumin (BSA) and casein. The results (**Figure S1**) suggest that albeit casein has a better blocking performance, it seems to mask the previous immobilized proteins, rendering them inaccessible. Conclusively, we employed BSA as blocking agent for subsequent experiments.

Since the PhyB/PIF interaction plays the key role in our experimental setup, we tested whether a competitor complex containing two PIF proteins shows better performance in terms of leakiness, i.e., fewer unspecific release during red-light illumination due to increased avidity. Therefore, the release of competitor complexes containing one or two PIF proteins was measured over a time range of 80 minutes. Additionally, to determine the unspecific release of the nitrocellulose membrane itself, the release of a biotinylated version of GFP-PIF6, immobilized directly on nAv, was determined. According to our findings (**Figure S2)**, the double PIF version has no advantages over the single version regarding the unspecific release (red light, 660 nm). In fact, the release of competitor molecules can be attributed to the unspecific release of the nitrocellulose substrate itself as the biotin-GFP-PIF6 version shows similar values than the two other non-released samples. However, far-red light illumination (740 nm) yields a higher release for the single PIF version, which is why we conducted all following experiments with this competitor version.

In order to find an assay architecture in terms of the assay area and their positioning, we tested three different variants regarding the diffusion time and fluid distribution on the distinct areas (**Figure S3**). For naked eye visualization, highly concentrated fluorescent protein mCherry (mCh) or the small molecule fluorescein was used. For both substances, the design variant where a small square shaped receiver area is enclosed by a larger square shaped sender area showed the best performance and was, therefore, employed for the following experiments.

For the proof-of-principle assay, we first demonstrated the light-induced competition of analyte and competitor. To simulate and verify the competitive mode of the OptoAssay, three conditions were tested: (i) red light illumination without analyte where no release of the competitor is expected, (ii) far-red light illumination without analyte where a release but no competition of analyte and competitor is possible, and (iii) far-red light illumination with analyte, where both a release and competition are expected. The fluorescence intensity measurements of the nitrocellulose membranes after illumination show a strong decrease in fluorescence signal of the sender membranes illuminated with far-red light compared to those illuminated with red light. Far-red light illuminated samples without analyte display a higher fluorescence in the receiver area than the ones where analyte was added (**Figure 2A**). This can also be confirmed by quantifying fluorescence intensities of the membranes (**Figure 2B**). Also, the measurements of the supernatant collected (**Figure 1**confirm this observation as there is a lower fluorescence intensity for the far-red light illuminated samples without analyte than analyte-containing samples. When comparing the leakiness (i.e., unspecific release of competitor during red light illumination) to the red-light induced specific release by dividing the fluorescence intensity of the far-red by the red light illuminated sample, there is a 6.1-times increase in signals of the sample with analyte but only a 4.5-times signal increase for the sample without analyte. This could be due to the competitor binding to the receiver area and, therefore, leaving less competitor in the supernatant. While there is a high fluorescence intensity obtained on the receiver from the sample treated with far-red light and without analyte, only a low fluorescence is detectable for the far-red light illuminated samples with analyte. A quantification of the fluorescence intensities of receiver membranes after illumination shows a 115% signal increase when comparing far-red light illuminated samples with red light illuminated ones without analyte and only a 22% increase in signals of far-red light illuminated membranes with analyte. This results in a fluorescent signal increase of 163% when comparing far-red light illuminated samples with and without analyte.

**Figure 2.**
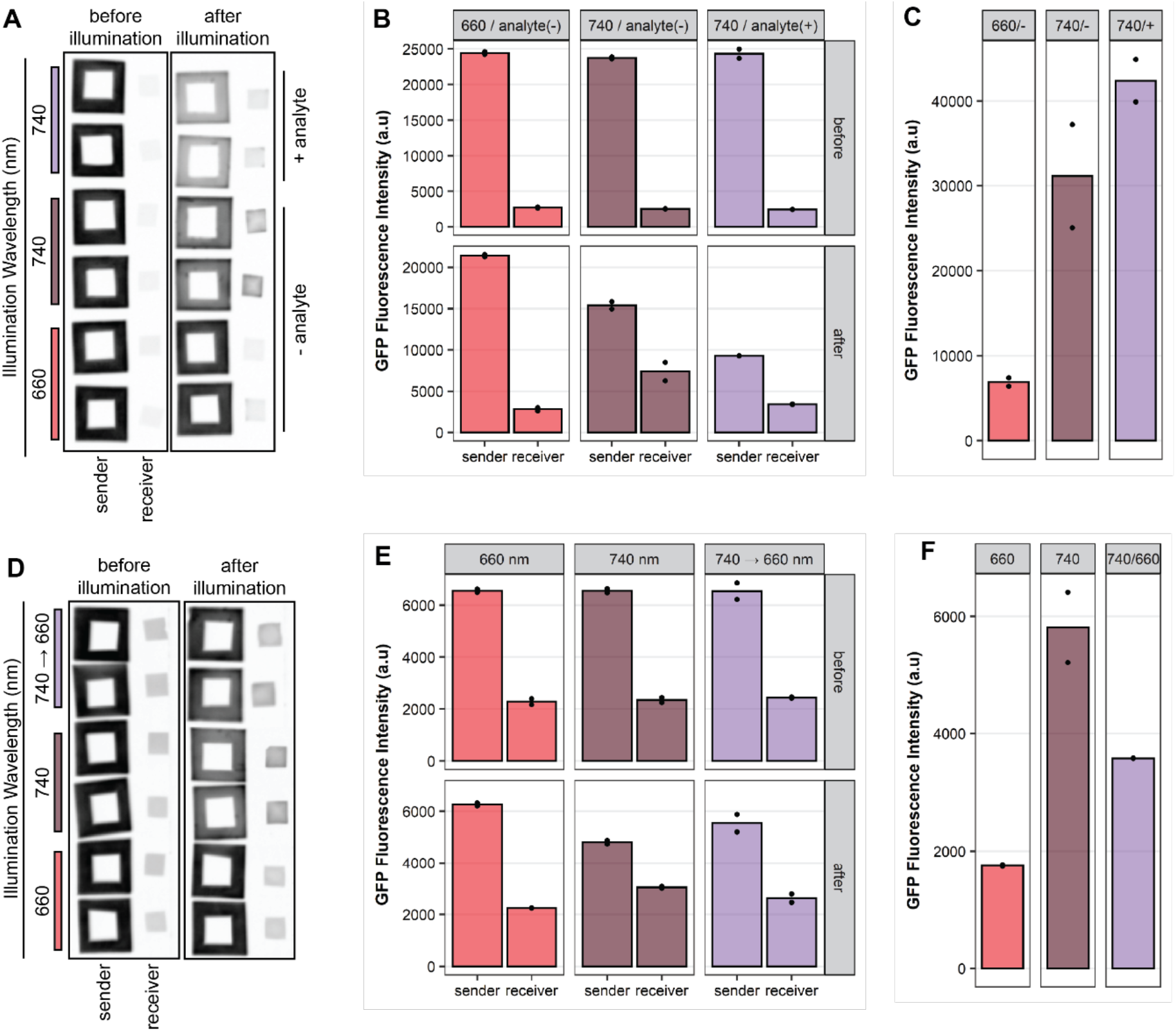
**A)** Release of the competitor and competition with an analyte on nitrocellulose membranes. Fluorescence images of nitrocellulose membranes before and after illumination. Sender membranes were incubated with nAv and then blocked with BSA. Next, biotinylated PhyB was immobilized. The competitor complex (his-GFP-PIF6, 30 mg ml^−1^) was bound to PhyB under red light illumination. The receiver membranes were coated with anti his-tag antibody and subsequently blocked with BSA. For the measurement, sender and receiver membranes were assembled and covered with buffer or buffer containing the his-tagged analyte protein which was added at the same time as illumination started. The samples were illuminated from above either with red or far-red light for 40 minutes. The membranes were washed under illumination with the corresponding light and subsequently imaged. **B)** Membrane fluorescence intensity values quantified from A in arbitrary units (a.u.) **C)** Supernatant fluorescence of the samples from A after illumination in a.u. **D)** Release and rebinding of the competitor on nitrocellulose membranes. Fluorescence images of nitrocellulose samples before and after illumination. Sender membranes were incubated with nAv followed by blocking with BSA and immobilization of biotinylated PhyB. The competitor complex (his-GFP-PIF6, 1.3 μg ml^−1^) was associated with PhyB under red light illumination. The receiver membranes were first coated with anti-his antibody (10 μg ml^−1^) and subsequently blocked. Sender and receiver areas were assembled, and buffer was added to cover both areas. Illumination from above for 40 minutes either with red or farred light or first 40 minutes with far-red and then 40 minutes with red light was carried out. Membranes were washed under illumination with the corresponding light and subsequently imaged. The contrast of the image was adjusted for better visualisation. The original images can be seen in **Figure S4**. **E)** Membrane fluorescence intensity values quantified from D in a.u. **F)** Supernatant fluorescence of the samples from D after illumination in a.u.

The last step of the OptoAssay comprises of the rebinding of the unbound competitor molecules back to PhyB on the sender membrane which could enable a wash-free signal readout. To achieve this, one sample was illuminated first with far-red light to release the competitor and then with red light to rebind it. Control samples that were either illuminated with red or far-red light only were included as controls. The fluorescence images of the membranes (**Figure 2D**) after illumination indicate that the control samples worked as anticipated. In the case of the sender membranes, the measured intensity of the far-red sample is lower than the red light one. Also, the binding of the competitor on the receiver membrane can be observed for the (far-red light illuminated) release samples (**Figure 2E**). The rebinding samples (illuminated with far-red and then red light) display a slight increase in fluorescence intensity of sender membranes. The rebinding of the unbound competitor was also demonstrated by measuring the supernatant fluorescence (**Figure 2F**) which corresponds the amount of free, unbound competitor. Here, the fluorescence intensity of the rebinding sample has only is only 61% compared to the release sample. This means that about 39% of the released competitor could be rebound to the sender membrane. When comparing the membrane intensities (**Figure 2E**) before and after illumination, a reduction of fluorescence of the leakiness sample (red light) of 5% and 2% for sender and receiver, respectively, was observed. The release sample shows 36% reduction of intensity on the sender and a 23% increase on the receiver membrane. In the case of the rebinding sample, however, there is only a reduction of only 18% on the sender membrane which indicates a partial rebinding. The receiver membranes prove only an increase of 7%. For this experiment, lower amounts of competitor were applied compared to the previous experiment to ensure a higher rebinding efficiency, as depicted in **Figure S4**. The general lower fluorescence intensity of the rebinding samples, however, could be attributed to the longer incubation time of 80 minutes compared with 40 minutes used for the other samples. The longer incubation period might have led to higher dissociation of PhyB and PIF6 or even the unspecific release of nAv from the nitrocellulose membrane (see **Figure S2**).

Having successfully demonstrated the proof-of-concept of the OptoAssay on a nitrocellulose substrate, we aimed to the test the general functionality of the optogenetic components on other substrates. As an alternative planar substrate, we used PMMA which is low-priced and thus convenient for mass production of a potential POC device. A preliminary experiment (**Figure S5**) showed that nAv directly absorbs to untreated PMMA which circumvents the need for a surface functionalization. Subsequently, we immobilized biotinylated PhyB on the nAv coated surface. With this setup, we could demonstrate a spatially resolved release of the competitor complex from a PMMA substrate (**Figure S6**). However, we did not pursue with PMMA as a substrate for the OptoAssay because the total amount of immobilized photoreceptor was lower compared to nitrocellulose.

The next substrate tested were functionalized beads that are very often used as solid phase materials in bioassay applications in clinical and POC diagnostics^14,15^. In this setting, we verified the functionality of the optogenetic system on a bead-based platform where nAv functionalized agarose beads are employed to immobilize the biotinylated PhyB. We simulated the reversibility of releasing and rebinding the PIF6 molecule as the competitor from PhyB as described in^10^, immobilized on agarose beads (**Figure S7**). We could again show a light-dependent release as well as partial rebinding of competitor molecules from and to the substrate suggesting the likely generic applicability of the OptoAssay in various assay formats.

## Conclusion and Outlook

Here, we have successfully demonstrated the proof-of-principle of an OptoAssay that enables, in contrast to conventional POC tests like LFAs, a bi-directional sample flow without any pump or flow control systems for a wash-free signal readout. Through a photoreceptor that can light-dependently interact with a binding partner, a competitor molecule can be released and later removed from the detection area by applying far-red and red light. Although other photoswitchable assays already exist^16,17^, they only allow for detection of specific molecules, whereas the OptoAssay can be universally applied by fusing the molecule of interest to the phytochrome B interaction partner PIF6. For example, the integrating of a z domain^18^, an IgG-binding domain, antibodies can be attached to the PIF6 construct and act as detection antibodies to add even more versatility.

The system could also be further expanded by using or combining different optogenetic switches that respond to distinct wavelengths. For instance, the blue light receptor cryptochrome 2 (Cry2) that forms heterodimers with its interaction partner cryptochrome-interacting basic helix-loop-helix (CIB1), or another system, light-induced dimer (iLID), which comprises of the blue light receptor light oxygen voltage 2 domain of *Avena sativa* phototropin 1-SsrA (AsLOV2) and its binding partner stringent starvation protein B (SspB) could be employed^19^. However, both photoreceptors only enable an active associating with its interaction partner, the dissociation is usually a slower passive event induced by the absence of blue light. Therefore, the OptoAssay workflow might need to be adapted accordingly in order to use different photoreceptors.

Using a 3D printed PhotoBox along with a smartphone, the OptoAssay introduced can be employed for on-site applications. Herein, the PhotoBox could be extended by an excitation light and emission filter that allows for GFP detection using a smartphone. Various low-cost and easy sensing approaches in combination with a smartphone for a fluorescence readout have already been implemented^20^. Finally, automated sample illumination and evaluation of the results via a smartphone app could be realized for user friendly operation.

Another issue that must be addressed is the overall operation time of the OptoAssay which is, at the moment, mainly limited by diffusion of the biomolecules employed. Herein, there are two important factors: the overall distance the molecules have to travel between the two areas (*i.e.*, detection and immobilization zones) and the speed at which the molecules move. To decrease the distance, smaller assay areas could be designed and fabricated by micro/nanofabrication techniques. For increasing the speed, external energy, like ultrasonication, could be also applied to the system.

The OptoAssay provides the basics for a novel and dynamic bioassay format (independent of assay type and biomolecules employed, such as antibodies, proteins, or nucleic acids) through its light dependent two-way switch that can extend already existing POC devices and could pave the way for new classes of diagnostic devices in future, extending the capabilities of these state-of-the-art tools.

## Acknowledgements

The authors would like to thank the Deutsche Forschungsgemeinschaft (DFG, German Research Foundation) for funding this work under grant number 404478562 and Germany’s Excellence Strategy CIBSS – EXC-2189 – Project ID: 390939984. We would also like to thank Felicia Wieland for designing and building the PhotoBox.

## Competing Interest

The authors declare no competing interests.

## Contributions

N.U and M.H contributed equally to this work. C.D and W.W conceived the project. N.U., C.D., M.H and W.W. designed, analysed, and interpreted experiments. N.U performed experiments. C.D, M.H and WW supervised the project. N.U wrote the manuscript with support from all other authors.

## Methods

### Plasmids and oligos

The plasmids generated and used in this study are described in **Table 1** along with the templates and oligos for the polymerase chain reaction (PCR) to generate plasmids using Gibson cloning^21^. The sequences of the oligos used are listed in **Table 2**.

**Table 1.**
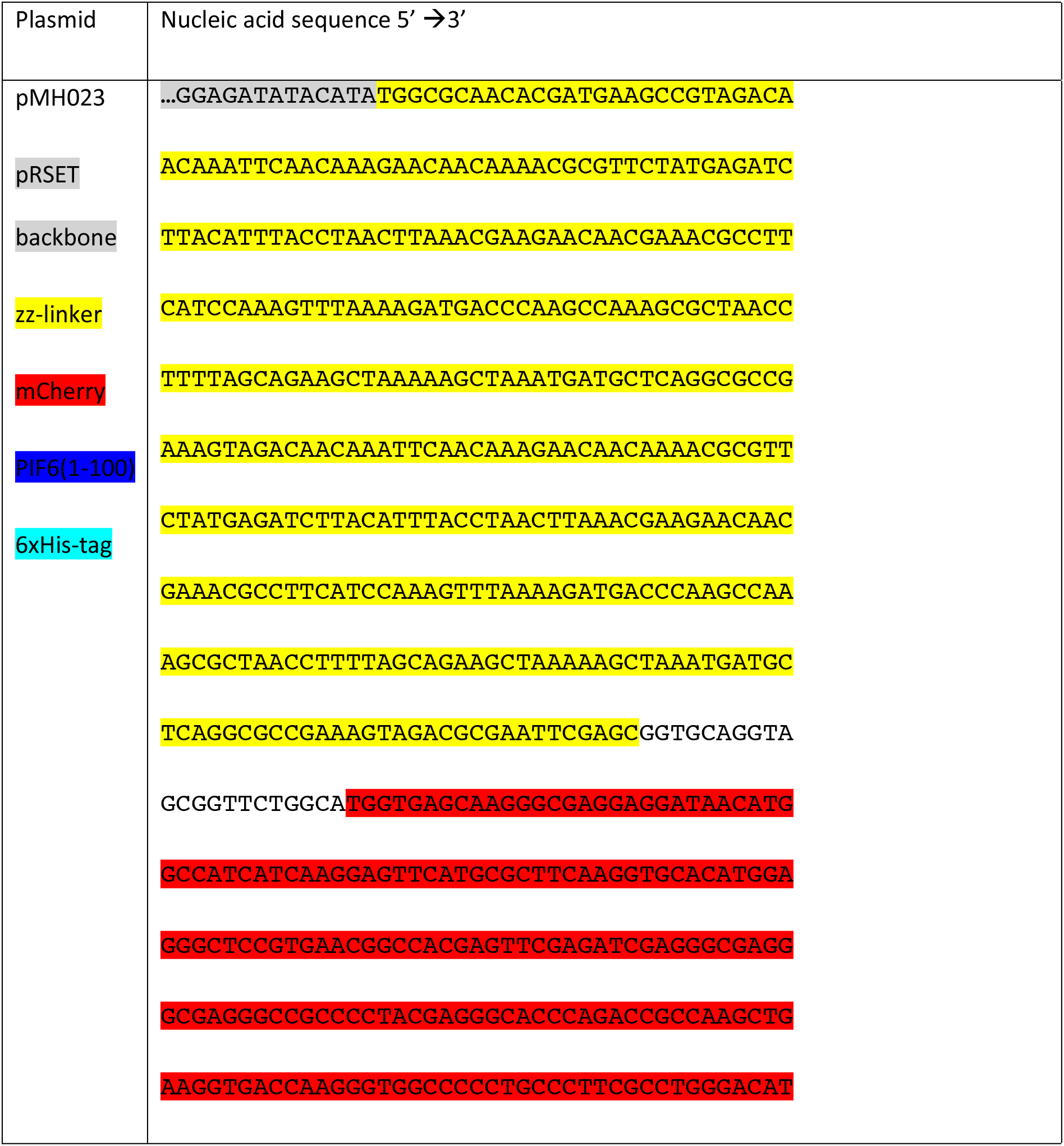
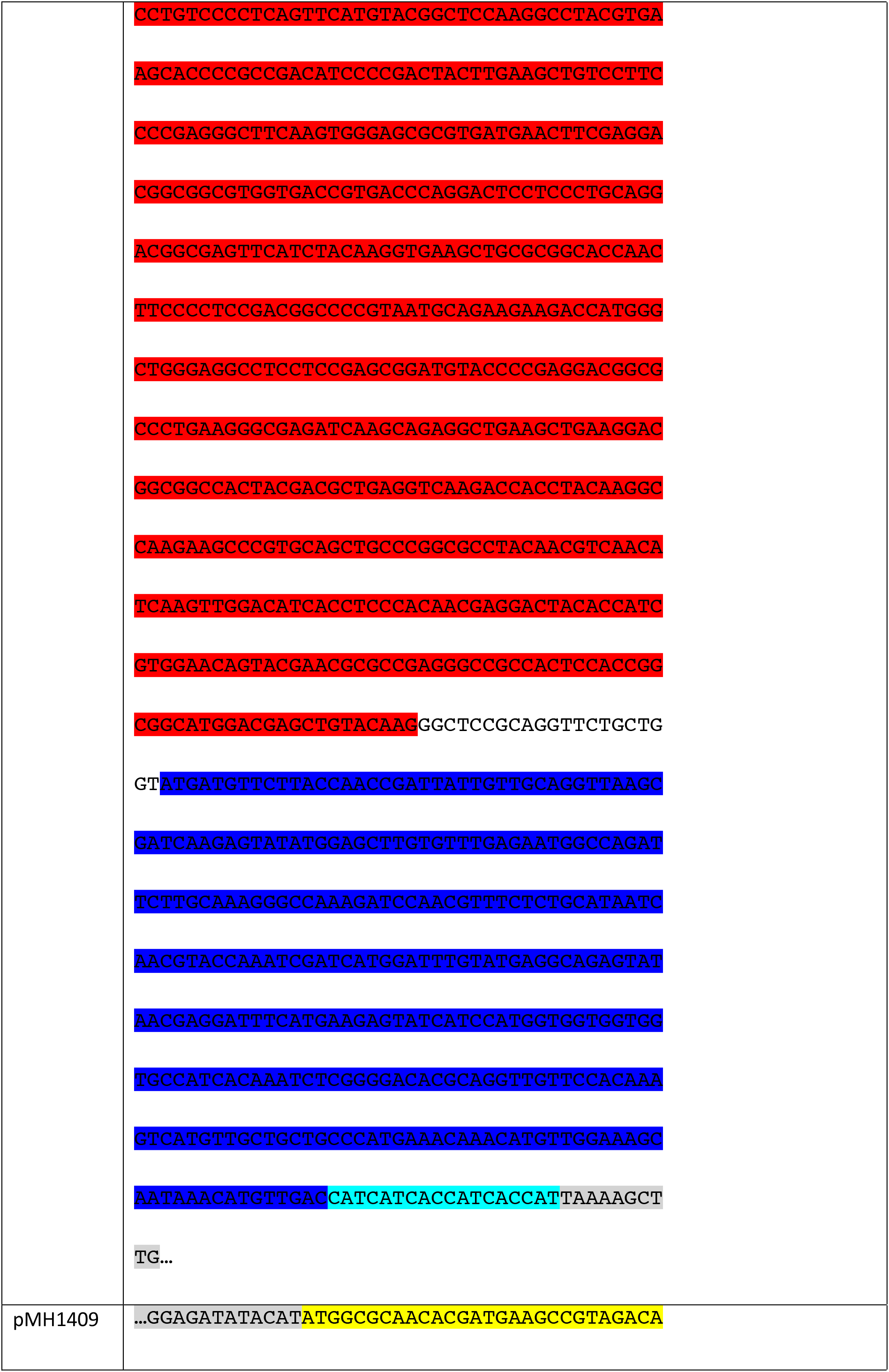
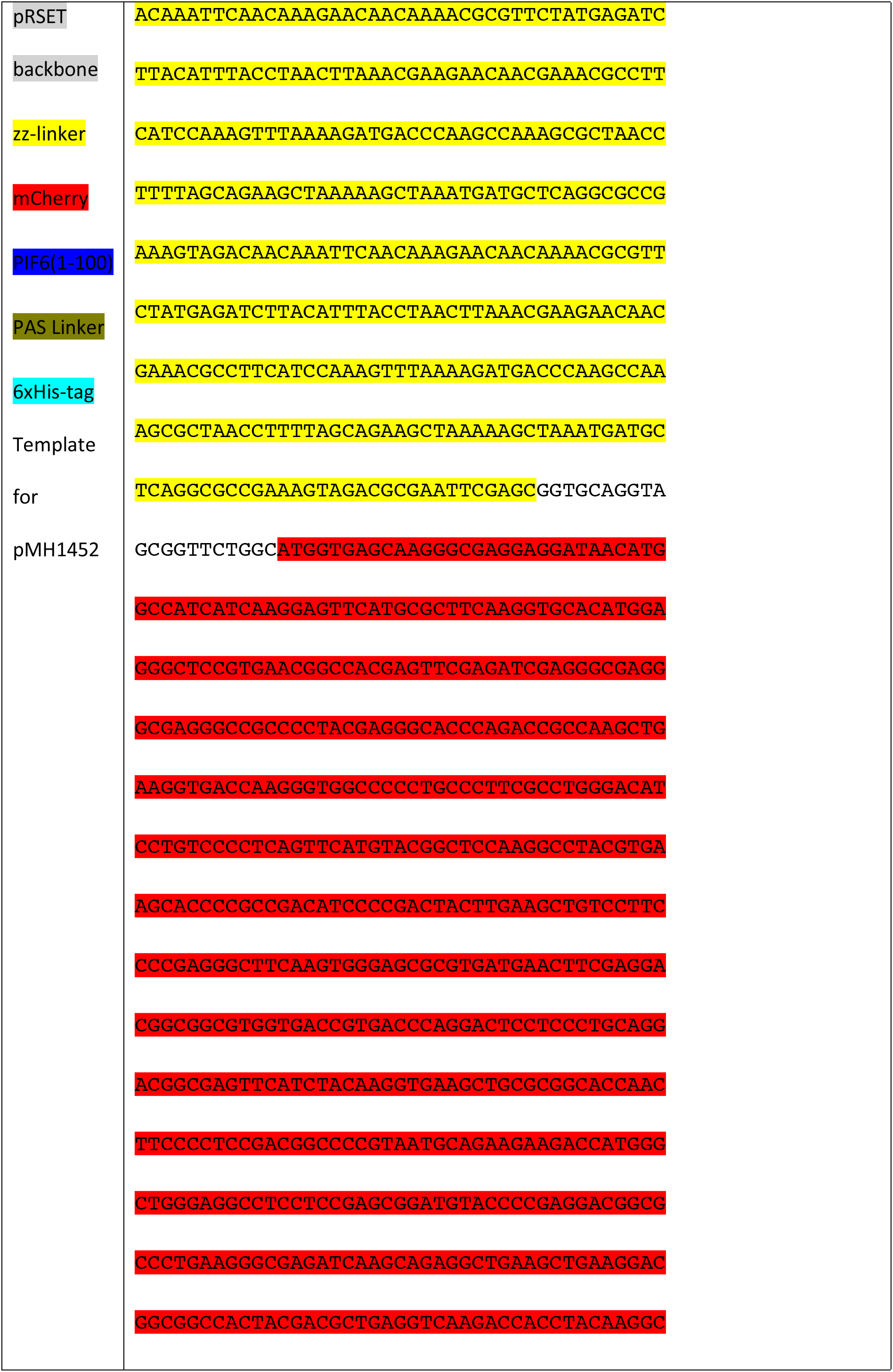
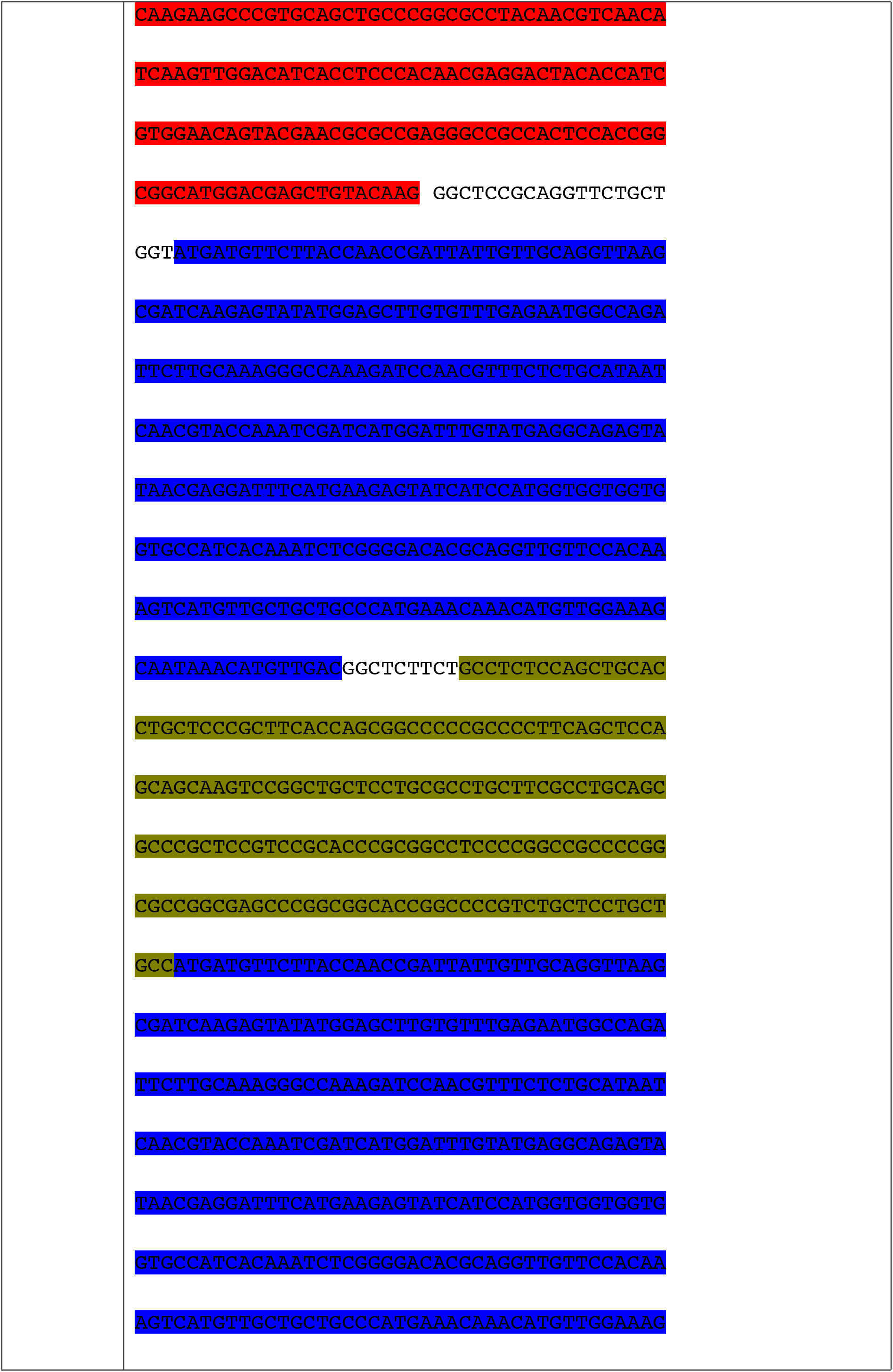
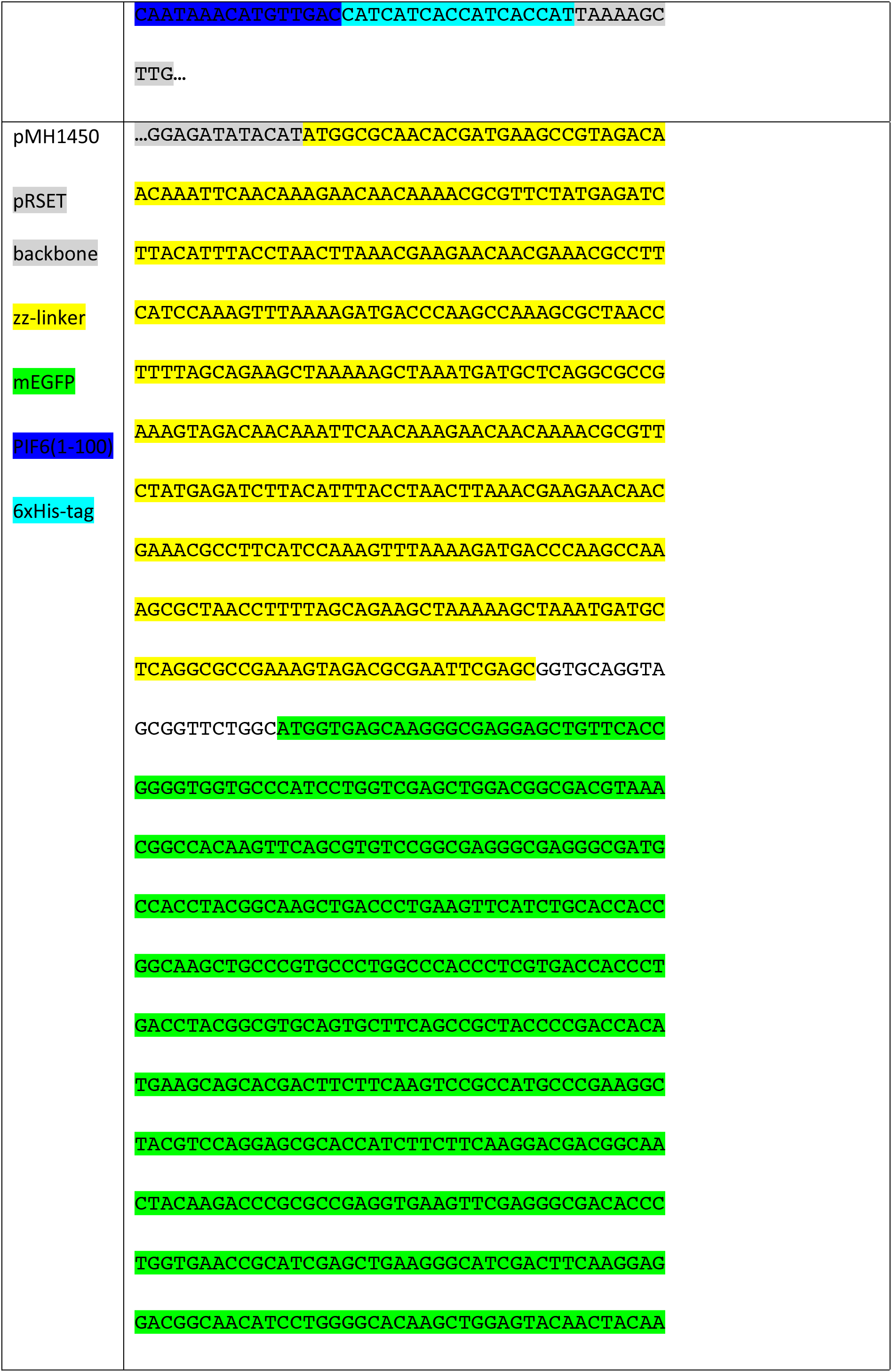
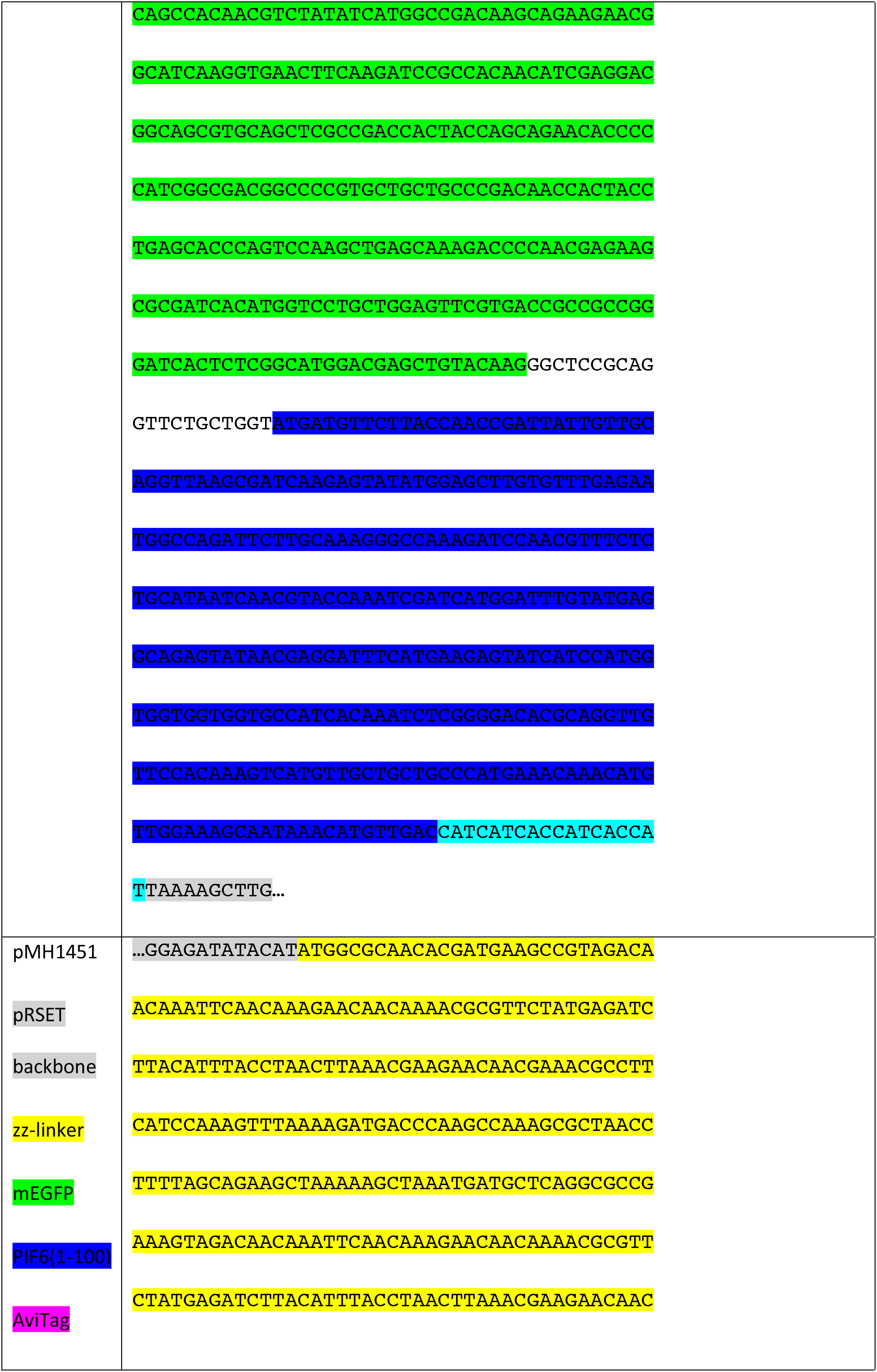
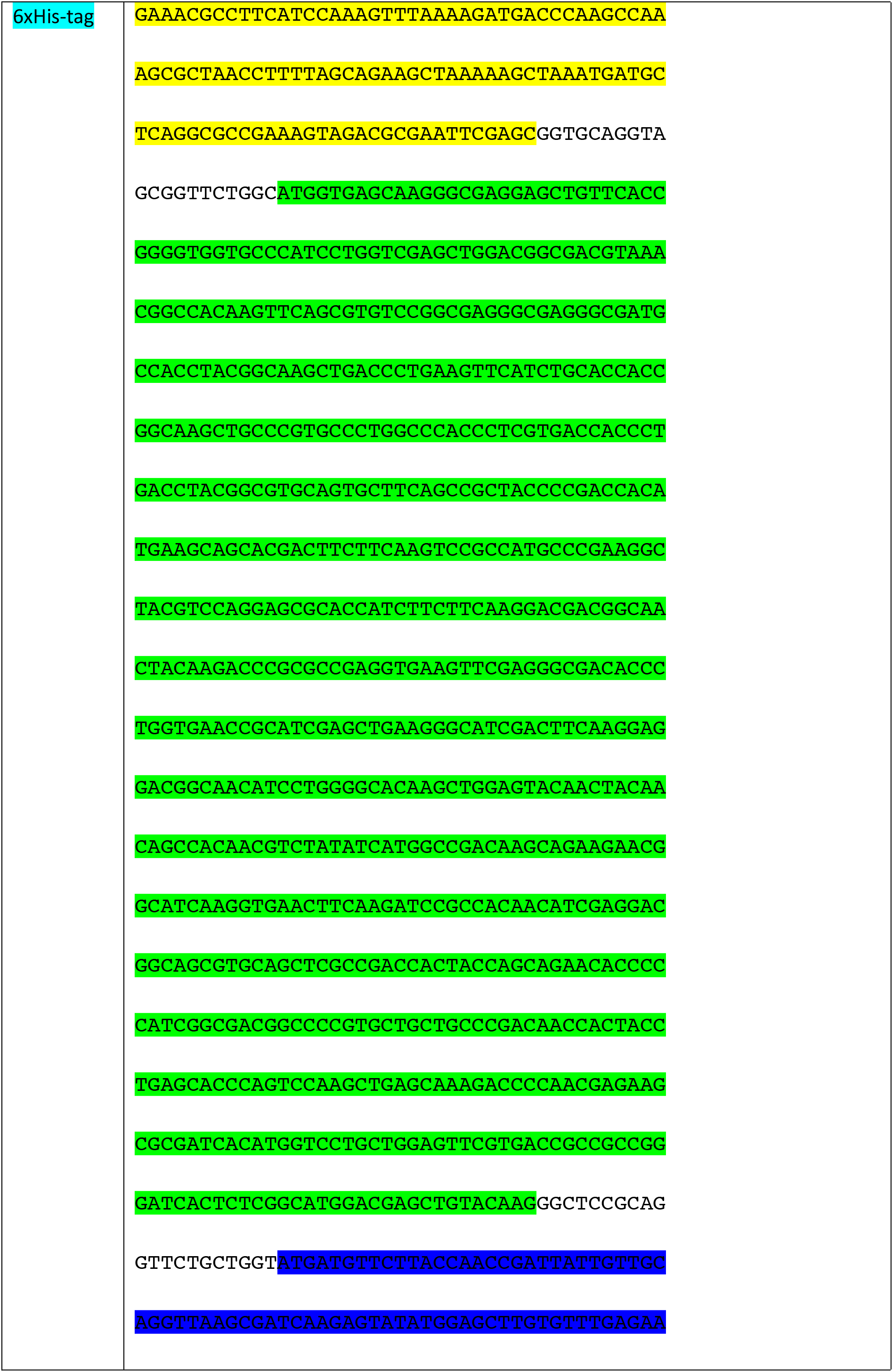
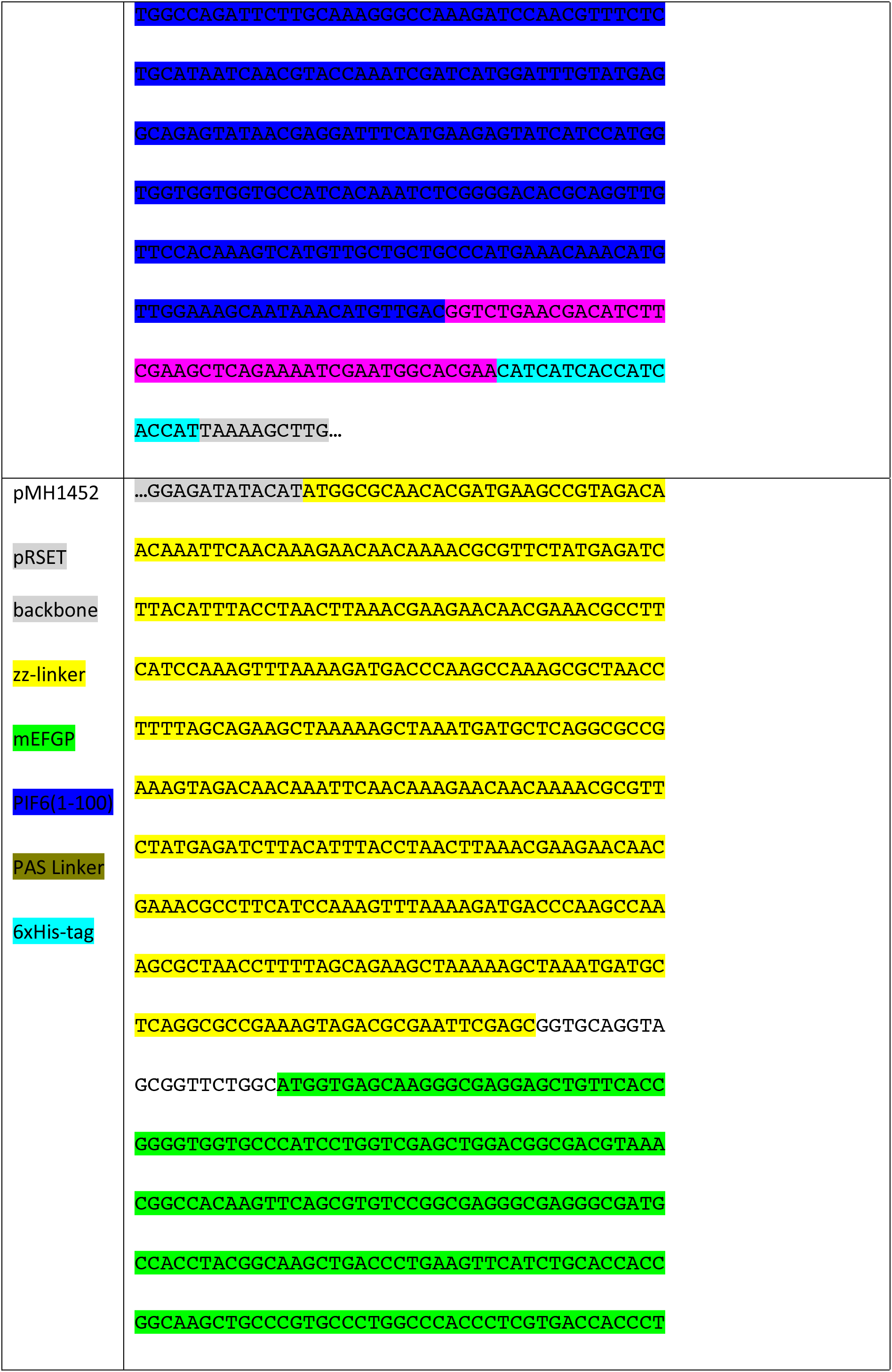
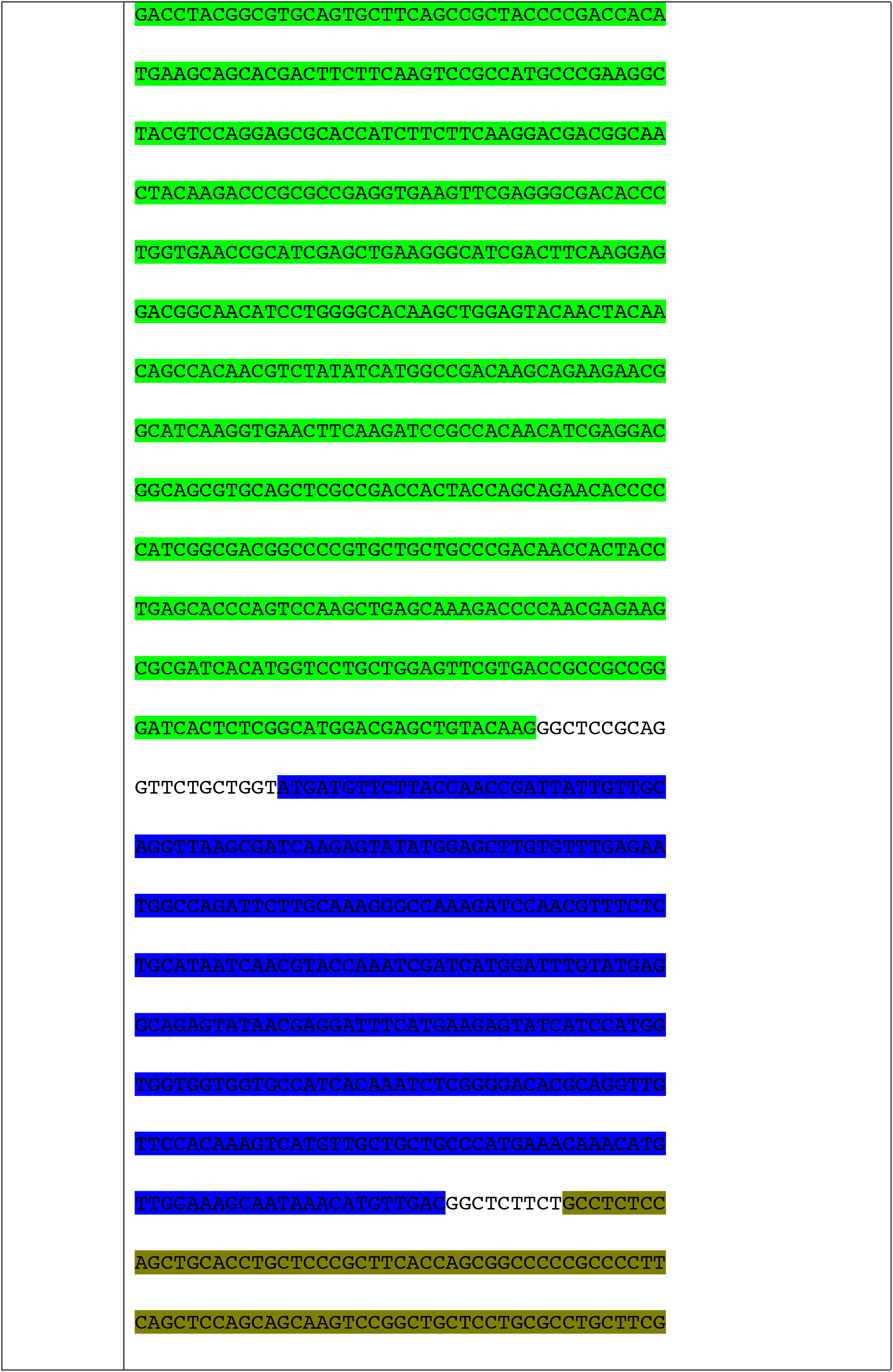
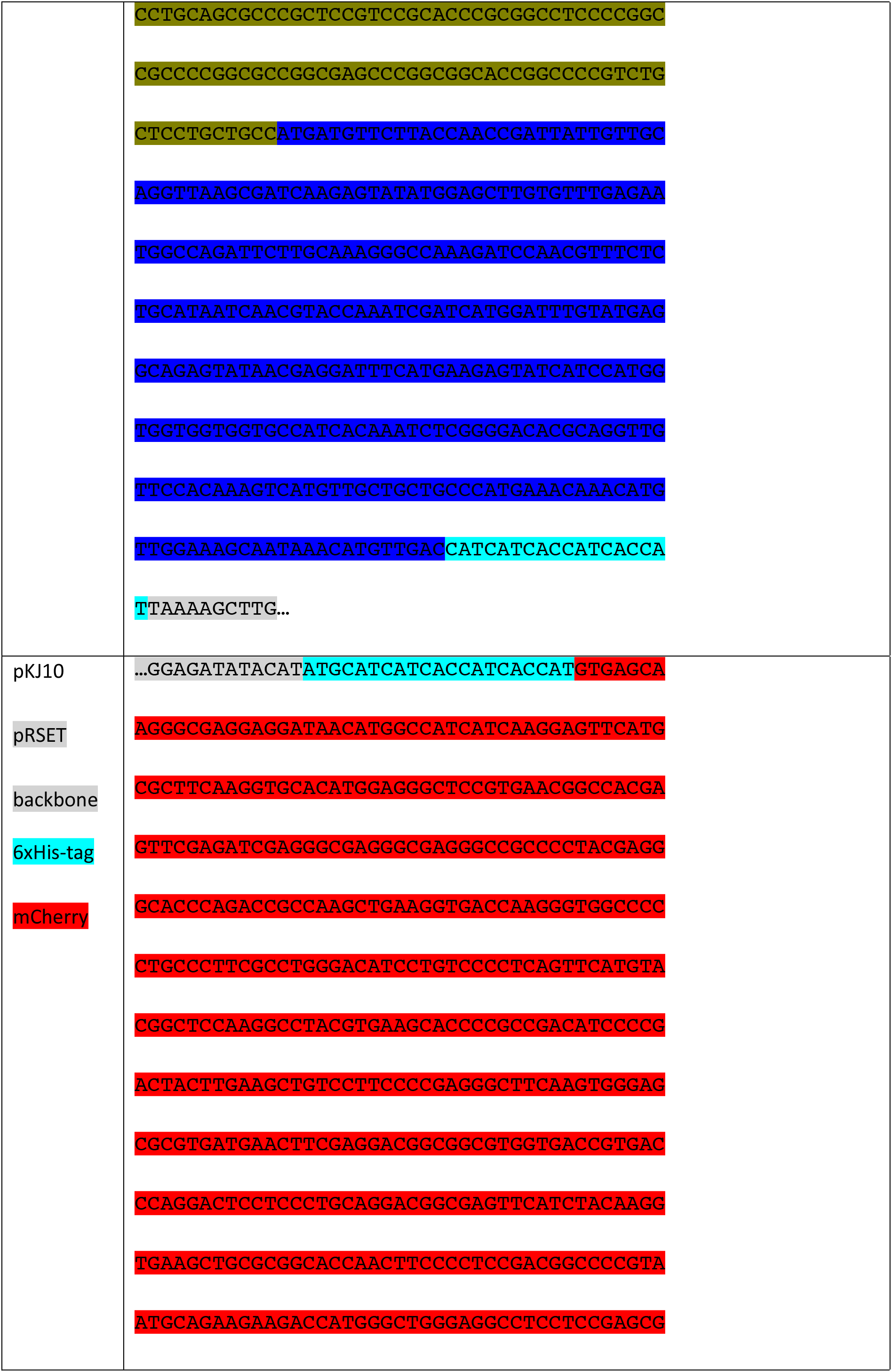
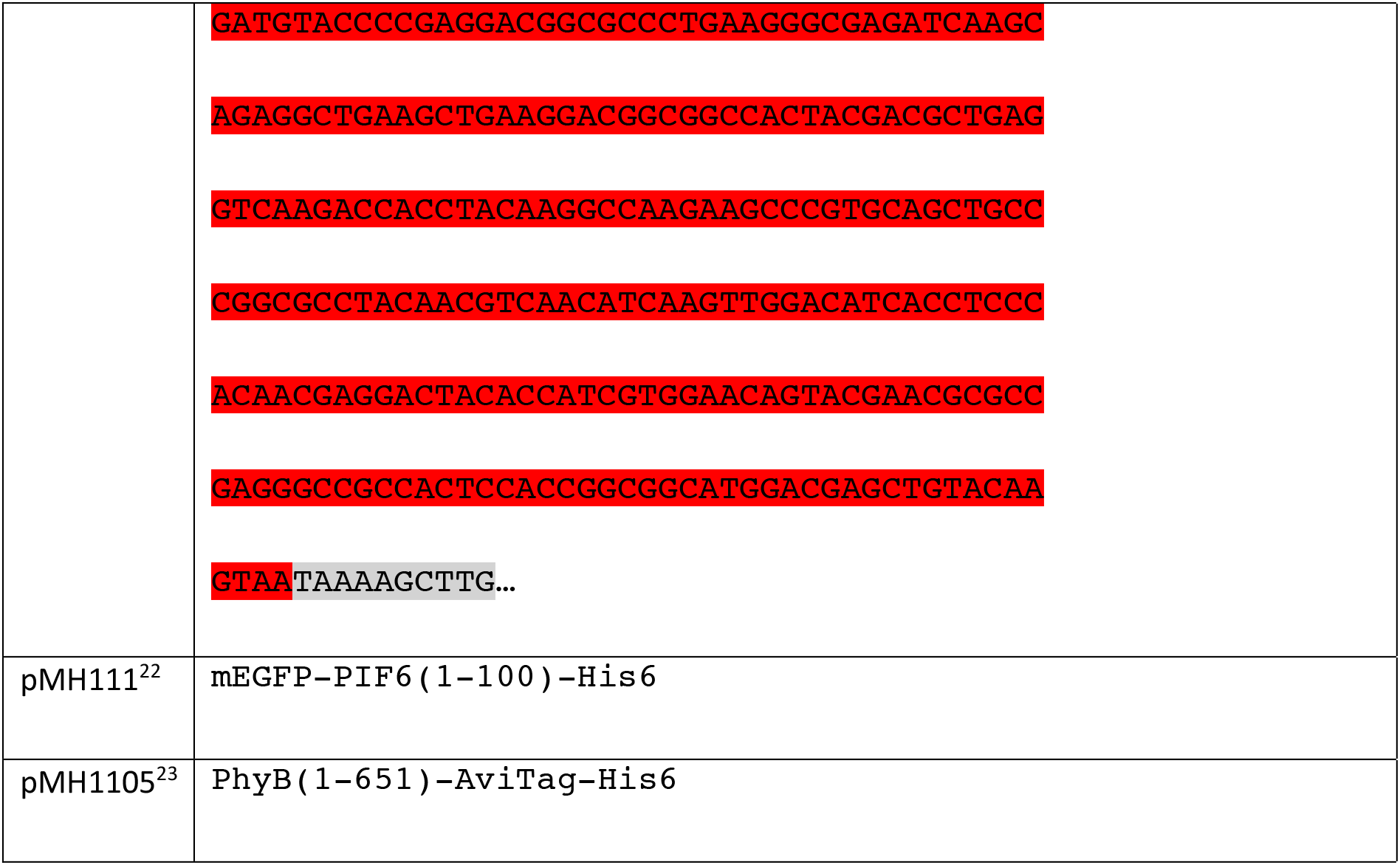
Nucleic acid sequence of plasmids used in this study

### Protein production and purification

#### PhyB constructs

PhyB-AviTag was produced by high-cell-density fermentation as described previously^5^. PhyB-AviTag (pMH1105) bacterial pellets, stored at –80 °C, were dissolved in Ni-Lysis buffer 50 mM NaH_2_PO_4_, 300 mM NaCl, 10 mM imidazole, pH 8.0),10 ml buffer per 1 g bacterial pellet. The suspension was lysed using a French press (APV 2000, APV Manufacturing) running three cycles at 1,000 bar. The lysate was centrifuged for 17,000 × g for 30 min at 4 °C, then the supernatant was transferred to a new vessel and centrifuged again for 30 min. The protein was purified using a Ni-NTA Superflow column (Qiagen, Netherlands, 30761) on a Äkta Express fast protein liquid chromatography system (FPLC, GE Healthcare, USA). The lysate was loaded on the column which was afterwards washed with 12 column volumes Ni-wash buffer (50 mM NaH_2_PO_4_, 300 mM NaCl, 20 mM imidazole, 0.5 mM tris(2-carboxyethyl)phosphine (TCEP), pH 8.0). The protein was eluted in 6 column volumes Ni-elution buffer (50 mM NaH_2_PO_4_, 300 mM NaCl, 250 mM imidazole, 0.5 mM TCEP, pH 8.0). The eluted protein was stored at 4 °C or frozen at –80 °C.

Before freezing of the protein, the protein was concentrated using centrifugal concentrators (Corning Inc., USA, 431491) and the buffer was exchanged to DPBS (2.7 mM KCl, 1.5 mM KH_2_PO_4_, 136.9 mM NaCl, 8.9 mM Na_2_HPO_4_ × 7H_2_O, pH 7.0) with a 10 mL dextran desalting column (ThermoFisher Scientific, USA, 431489). The protein was aliquoted, shock frozen in liquid nitrogen and stored at –80 °C.

#### PIF6 constructs

For protein production, the corresponding plasmid (table xx) was transformed into chemically competent E. coli BL21 Star (DE3) (Thermo Fisher Scientific, USA, C601003). The bacteria were cultured in LB medium supplemented with ampicillin (100 μg mL^−1^) to select the correctly transformed bacteria and were grown overnight at 37°C and 150 rpm. The overnight culture was transferred to new LB medium supplemented with ampicillin to reach an OD600 (optical density at 600 nm) of 0.15 and cultured at 30°C and 150 rpm. Upon reaching an OD600 of 0.5 – 0.8, the protein expression was induced using 1mM isopropyl-β-d-thiogalactopyranoside (IPTG) and transferred to 18°C incubation for 20h and 150 rpm. Cells were subsequently harvested by centrifugation at 6,500 g for 10 min and resuspended in lysis buffer and disrupted using a French Press at 1,000 bar or by sonication (Bandelin Sonopuls HD3100 (Bandelin, Germany). Lysates were clarified by centrifugation at 30,000 g for 30 min and loaded on a Ni-NTA Superflow column (Qiagen, Netherlands, 30761). After washing with each 20 column volumes of lysis and wash buffer (50 mM NaH_2_PO_4_, 300 mM NaCl, 20 mM imidazole, pH 8.0), purified proteins were eluted in 5 column volumes of Ni elution buffer (50 mM NaH_2_PO_4_, 300 mM NaCl, 250 mM imidazole, pH 8.0). Finally, the proteins produced were quantified by measuring the fluorescence or by Bradford assay (Bio-Rad, USA, 5000006).

### Illumination Experiments

#### OptoAssay

For sender membrane preparation the nitrocellulose membranes (ThermoFisher Scientific, USA, 88018) were cut into squares (8 × 8 mm) and put into a 24-well plate. The squares were incubated with 300 μL nAv (Thermo Fisher, USA, A2666) (1.5 mg mL^−1^ in DPBS) for 30 min. After washing with DPBS, the membranes were blocked with 300 μL blocking buffer (5% (w/v) BSA (PAN biotech, Germany, P06-139210) in DPBS)) for 1 h. Next, the membranes were incubated with 300 μL of biotinylated PhyB (pMH1105, 1 mg mL^−1^ in DPBS) for 30 min in the dark. After washing (0.1% BSA in DPBS), 300 μL of GFP-PIF6 (pHB111, 3 mg mL^−1^ for competition and 1.5 μg mL^−1^ for rebinding in blocking buffer) was incubated for 10 min under 660 nm illumination (30 s at 46 μmol m^−2^ s^−1^, afterwards at 15 μmol m^−2^ s^−1^) from above. The samples were afterwards washed again during 660 nm illumination.

Three sender area squares (3 × 3 mm) were put into a 24-well containing 300 μL of anti-His antibody (Merck, Germany, NOVG70796-3) (10 μg mL^−1^ in DPBS) and incubated for 30 min. The squares where then washed 300 μL washing buffer (0.1% BSA in DPBS) followed by a blocking step with 300 μL blocking buffer for 1 h.For the experiment, the sender membranes were assembled on parafilm, 80 μL of buffer (0.1% BSA in DPBS) was added and the samples were incubated in a humidified atmosphere. For competition, 80 μL of a his-tagged protein pyruvate dehydrogenase complex repressor (pdhR) (0.3 mmol L^−1^ in 0.1% BSA in DPBS) was used as analyte and was added at the same time as the illumination started. The samples were either illuminated with 660 nm (40 min at 15 μmol m^−2^ s^−1^) or 740 nm (10 min at 380 μmol m^−2^ s^−1^, 30 min at 200 μmol m^−2^ s^−1^) from above. Before visualization, the samples were rinsed with DPBS under illumination of the respective wavelength and put into a new well containing DPBS. The GFP fluorescence of the nitrocellulose membranes was imaged using an ImageQuant LAS 4000mini system (GE Healthcare, USA). For fluorescence measurements of the supernatant, 50 μL of the supernatant was collected after removing the nitrocellulose membrane, and subsequently measured using a microtiter plate reader Tecan infinite M200pro (Tecan Group, Switzerland) (excitation wavelength: 488 nm, emission wavelength: 530 nm); background fluorescence of the buffer (0.1% BSA in DPBS) was subtracted from each value before plotting.

#### Homogeneous or spatially resolved release from nitrocellulose

Nitrocellulose membranes (ThermoFisher Scientific, USA, 88018) were cut into squares (8 × 8 mm) and put into a 24-well plate. The samples were incubated 300 μL nAv (1.5 mg ml^−1^ in DPBS) for 30 min. After washing with DPBS for 2 min, the membranes were blocked with 300 μL BSA solution (1% in DPBS) for 30 min if not stated otherwise. Then, 300 μL of biotinylated PhyB (pMH1105, 1 mg mL^−1^ in DPBS or Ni elution buffer) was added and incubated for 30 min in the dark. After washing (0.1% BSA in DPBS), 300 μL of GFP-PIF6 (pMH1450, 3 × 10^−2^ mg mL^−1^ in 1 or 5% BSA dissolved in DPBS) was incubated for 30 min under 660 nm illumination (46 μmol m^−2^ s^−1^; alternating 1 min on, 5 min off) from above. The samples were afterwards washed during 660 nm illumination.

For spatially resolved release with a photomask, samples were put directly on the photomask, covered with buffer (0.1% BSA in DPBS) and illuminated from below with 740 nm (77 μmol m^−2^ s^−1^) light for 2 min. Immediately after illumination, the samples were rinsed with DPBS and put into a new well containing DPBS. Experiments without a photo mask were conducted in a 24-well plate, where a volume of 300 μL buffer was added before 740 nm (380 μmol m^−2^ s^−1^) or 660 nm (46 μmol m^−2^ s^−1^) illumination. Supernatant samples of 50 μL were taken at the indicated timepoints. Before visualization, the samples were with DPBS under illumination of the respective wavelength and put into new well containing DPBS.

For testing the different PIF6 versions (GFP-PIF6 (pMH1450) and GFP-2xPIF6 (pMH1452)) the nitrocellulose membranes where prepared as described above. For the samples treated with biotin-GFP-PIF6, the PIF construct was added directly to nAv functionalized and blocked membranes; this sample was only illuminated with red light. Fractions of the supernatant (50 μL) of all samples were taken at five time-points during the 80-minute illumination period. Fluorescence of the samples was measured using a plate-reader (excitation wavelength: 488 nm, emission wavelength: 530 nm); background fluorescence of the buffer (0.1% BSA in DPBS) was subtracted from each value before plotting.

#### Agarose Beads

For ten samples, 100 μL nAv functionalized agarose beads (ThermoFisher Scientific, USA, 29202), were washed with buffer (0.1% (w/v) BSA in DPBS) by centrifugation at 500 × g for 2 min. Then 800 μL PhyB (pMH1105, 2 mg mL^−1^ in Ni elution buffer) was added and incubated for 30 min in the dark. PhyB solution was removed, and the samples were washed. 600 μL of mCh-PIF6 (pMH23, 2.9 or 0.029 mg ml^−1^ in 0.1% BSA in DPBS) were added and incubated for 30 min under 660 nm illumination (46 μmol m^−2^ s^−1^; alternating 1 min on, 5 min off). After washing, the beads were re-suspended in 2 mL buffer (0.1% BSA in DPBS). For each sample, 200 μL of the re-suspended beads was transferred into a 1.5 mL reaction tube.

The reaction tubes were illuminated for 1 h at 660 nm (15 μmol m^−2^ s^−1^) or at 740 nm (380 μmol m^−2^ s^−1^) from above. After illumination, the samples were spun down at 500 × g for 2 min and 50 μL of the supernatant was taken for fluorescence measurements. The mCh fluorescence (excitation wavelength: 587 nm, emission wavelength: 625 nm) of the supernatant was detected using a microtiter plate-reader Tecan infinite M200pro; the background signal of the buffer was subtracted from each value before plotting.

#### Spatially resolved release from PMMA

PMMA (Röhm GmbH, Germany, 99524 GT, 1.175 mm) rectangles (2 × 1.5 cm^2^) were cut and put into a 6-well plate. The samples were incubated with 1 mL of nAv (ThermoFisher Scientific, USA, 31050) solution (1.5 mg ml^−1^ in DPBS) for 30 min. After washing with buffer (0.1% BSA in DPBS) for 2 min, PhyB (pMH1105, 2 mg mL^−1^ in Ni elution buffer) was incubated for 30 min in the dark. After washing, 1 mL of GFP-PIF6 (pMH1450, 1 mg ml^−1^ in Ni elution buffer) was incubated under 660 nm (46 μmol m^−2^ s^−1^; alternating 1 min on, 5 min off) illumination. The samples were afterwards washed again. Samples were put directly on the photomask, covered with buffer (0.1% BSA in DPBS) and illuminated from below with 740 nm (77 μmol m^−2^ s^−1^) for 1 min. Immediately after illumination, the samples were rinsed with DPBS and put into a new well containing DPBS. For visualization, the GFP fluorescence of the PMMA slides was imaged using the ImageQuant LAS 4000mini system.

### Assay Design Diffusion experiment

Cellulose sender areas were incubated in solution of a fluorescent protein mCh (pKJ10, molecular weight: 27,545 g mol^−1^, concentration: 0.2 mM) or fluorescein (molecular weight: 332 g mol^−1^, concentration: 10 mM) for 10 minutes. Afterwards, the treated sender and the untreated receiver membrane were assembled. Sender and receiver membranes were covered with DPBS, then images of the membranes were taken immediately after assembly and after 1, 5, 15, and 30 minutes.

### PhotoBox with integrated electronics

The 3D printed PhotoBox^24^ serves as incubation unit by illumination with 660 nm (12 μmol m^−2^ s^−1^) and 740 nm (185 μmol m^−2^ s^−1^) light and as imaging box with a smartphone-based readout. For smartphone, a Samsung Galaxy A5 (2017) (Samsung Electronics, South Korea) is used. The images are taken during flashlight illumination with the integrated smartphone camera. The LEDs can be switched on and off via an Arduino nano, which receives the signal via an HC-05 Bluetooth module. The Arduino nano is coded, so that the user only needs to send either “660 nm” or “740 nm” via smartphone to the HC-05 in order to turn on/off the LEDs. Here, a standard application for Android is used as Bluetooth terminal to connect to HC-05.

The PhotoBox was designed with SolidWorks 2014 (Dassault Systemes SolidWorks Corp., France) and printed out of black acrylonitrile butadiene styrene (ABS) with the Ultimaker 3 (Ultimaker B.V., Netherlands). The PCB was designed with EAGLE software and produced by the company Beta LAYOUT Ltd. (Germany) and the soldering of the electronic components was done in-house.

### Data analysis

The images were analyzed using the software ImageJ (University of Wisconsin-Madison, USA). The mean intensity values of the whole nitrocellulose membranes/PMMA pieces for the illuminated and not illuminated areas were analyzed with the area selection tool. In order to determine the final value of intensity, the mean background intensity was subtracted, and the obtained values were normalized. For normalization, intensity values of plain buffer were subtracted from each value.

---

**Figure S1.**
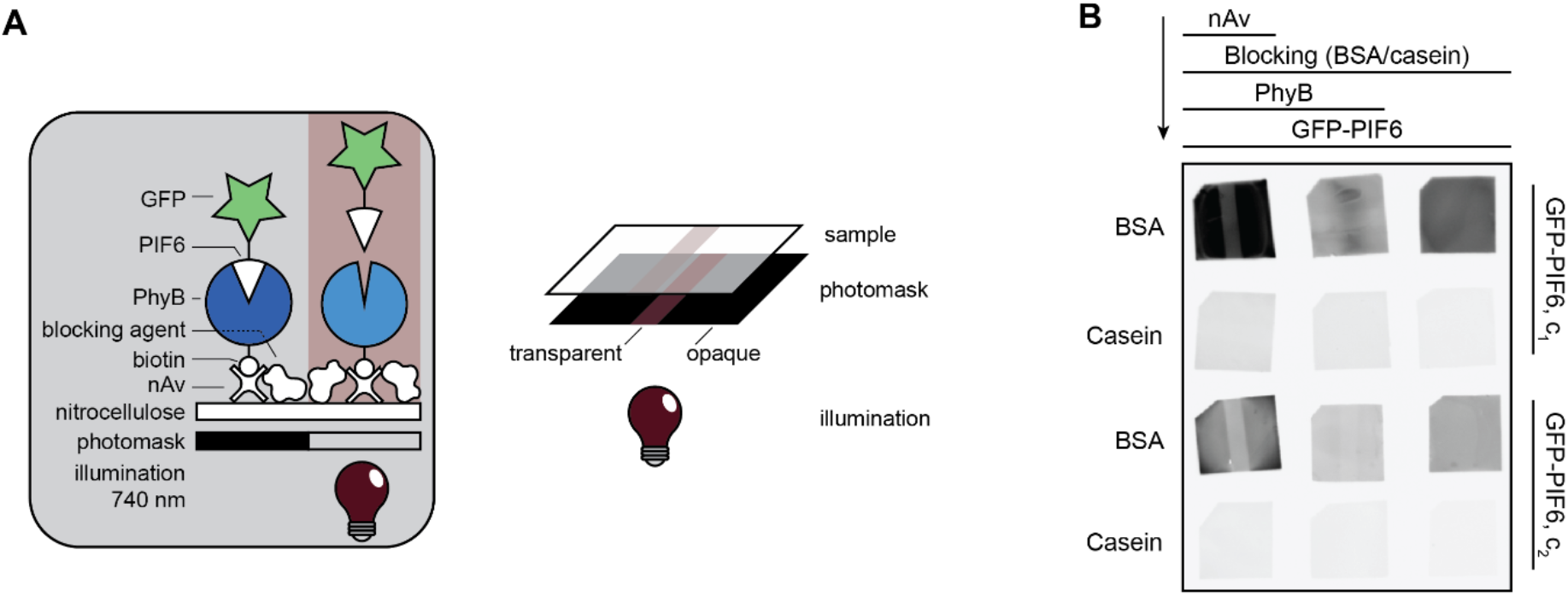
Blocking Strategy. **A)** Experimental setup. The nitrocellulose membrane was incubated with nAv, then blocked with a blocking agent (BSA or casein). After addition of PhyB, GFP-PIF6 was immobilized on PhyB during red light (660 nm) illumination and finally released through far-red light (740 nm) illumination. The scheme shows how the sample is luminated trough the transparent part of the photomask. **B)** GFP fluorescence of nitrocellulose membranes after the addition of GFP-PIF6 and illumination with far-red light (740 nm) through a photomask for 2 min. The samples were incubated with two different concentrations of GFP-PIF6 (c_1_ = 3 × 10^−2^ mg mL^−1^ and c_2_ = 0.6 × 10^−2^ mg mL^−1^) during red light illumination. The samples were covered with buffer before illumination with far-red light through a photomask.

**Figure S2.**
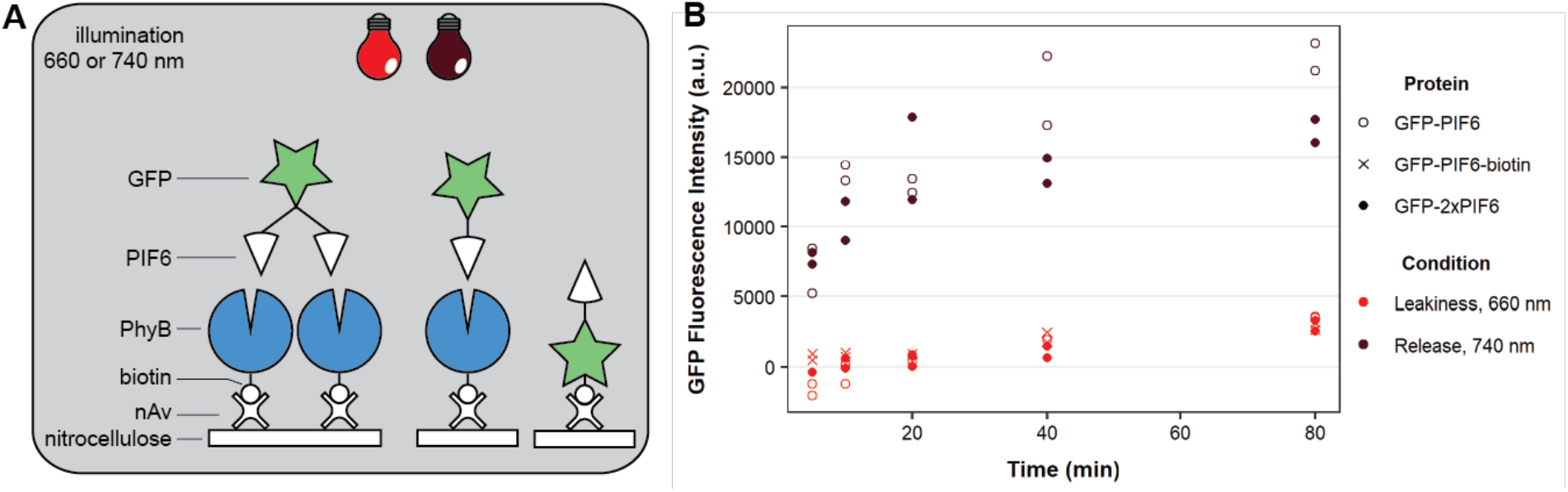
Different PIF6 Versions. **A)** Experimental setup. Neutravidin (nAv) treated nitrocellulose membranes were either incubated with PhyB before addition of GFP-PIF6 and GFP-2xPIF6, respectively, or GFP-PIF6-biotin was directly immobilized on the nAv-coated membranes. The samples were then illuminated with red (660 nm) or far-red (740 nm). **B)** Release kinetics of the model cargo GFP from functionalized nitrocellulose. The samples containing different cargo proteins GFP-PIF6, GFP-PIF6-biotin or GFP-2×PIF6 (3 × 10^−2^ mg mL^−1^) were illuminated from above in a 24-well plate, covered with buffer, for 80 min either with red (5 μmol m^−2^ s^−1^) or far-red light (380 μmol m^−2^ s^−1^). Graph shows the GFP fluorescence of supernatant over a time range of 80 min in a.u. 50 μL of the supernatant at timepoints 5, 10, 40 and 80 min were taken for GFP fluorescence measurements. The background fluorescence of the buffer was subtracted from each value. Red markings represent samples illuminated with red light, while dark-red ones the samples illuminated with far-red light.

**Figure S3.**
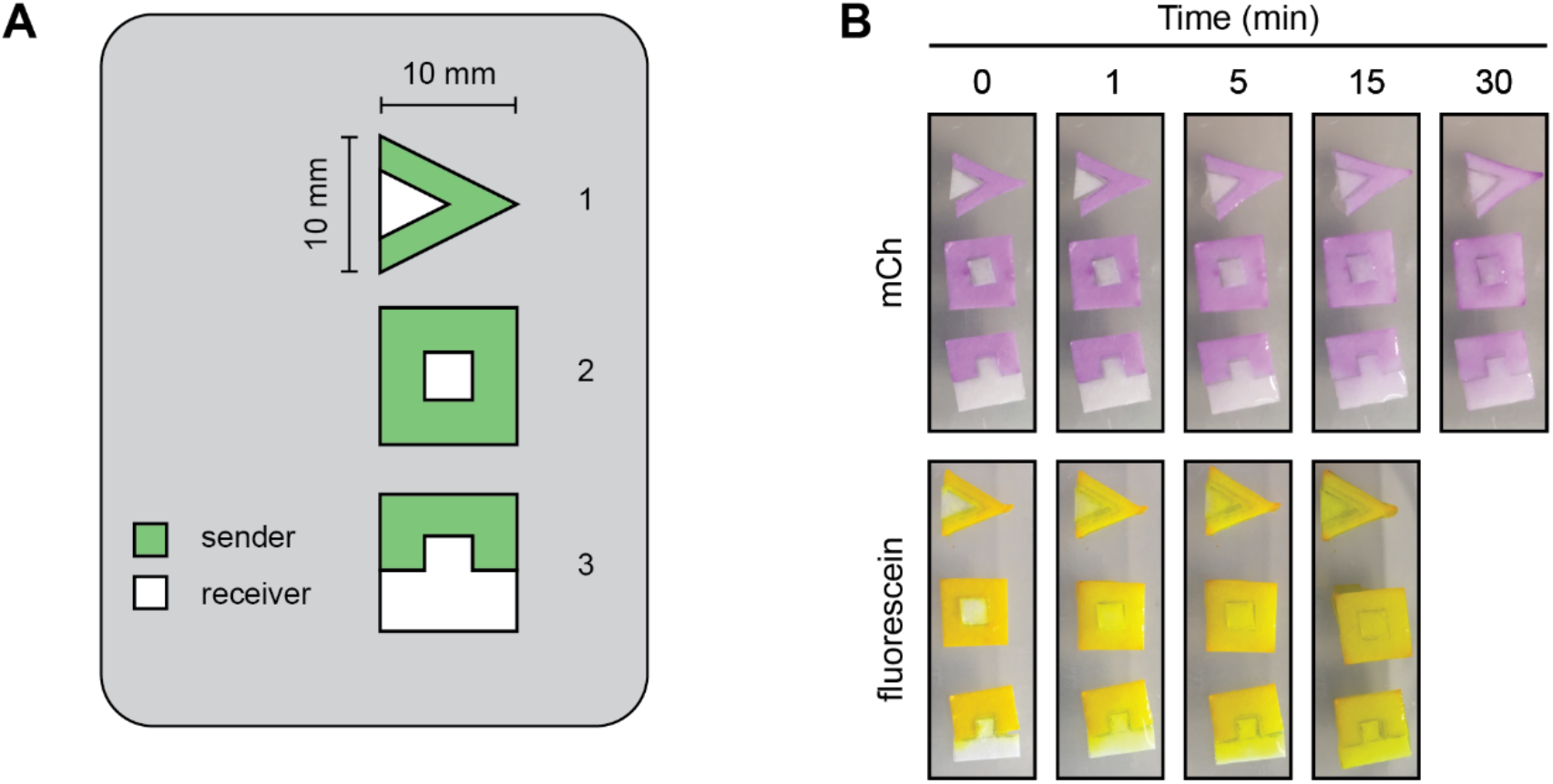
Geometry of OptoAssay Areas. **A)** Design of sender and receiver area. Sketch of different design variants. **B)** Images of the diffusion measurement from sender to receiver membrane. Cellulose sender areas were incubated in solution of the fluorescent protein mCherry (mCh) or the small molecule fluorescein. The treated sender and the untreated receiver membranes were assembled and covered with buffer. Images were taken immediately after assembly and after 1, 5, 15 and 30 minutes.

**Figure S4.**
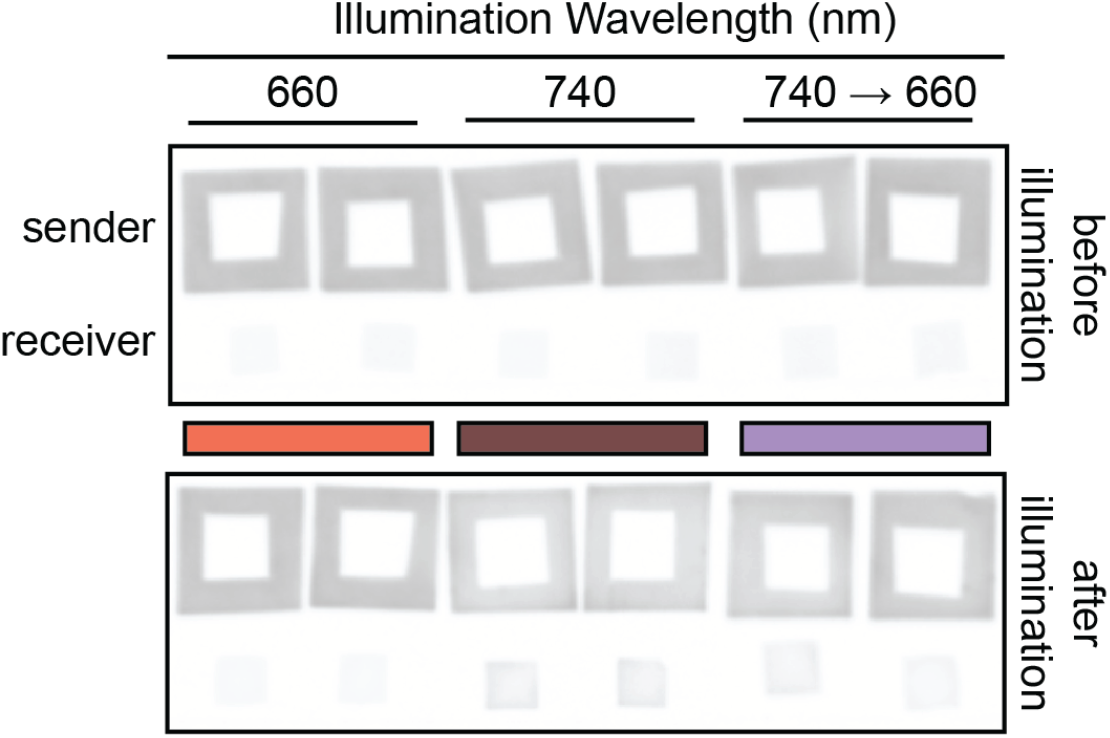
Original Membrane Intensities. Membrane intensities of Figure 2D with the same contrast settings as Figure 2A.

**Figure S5.**
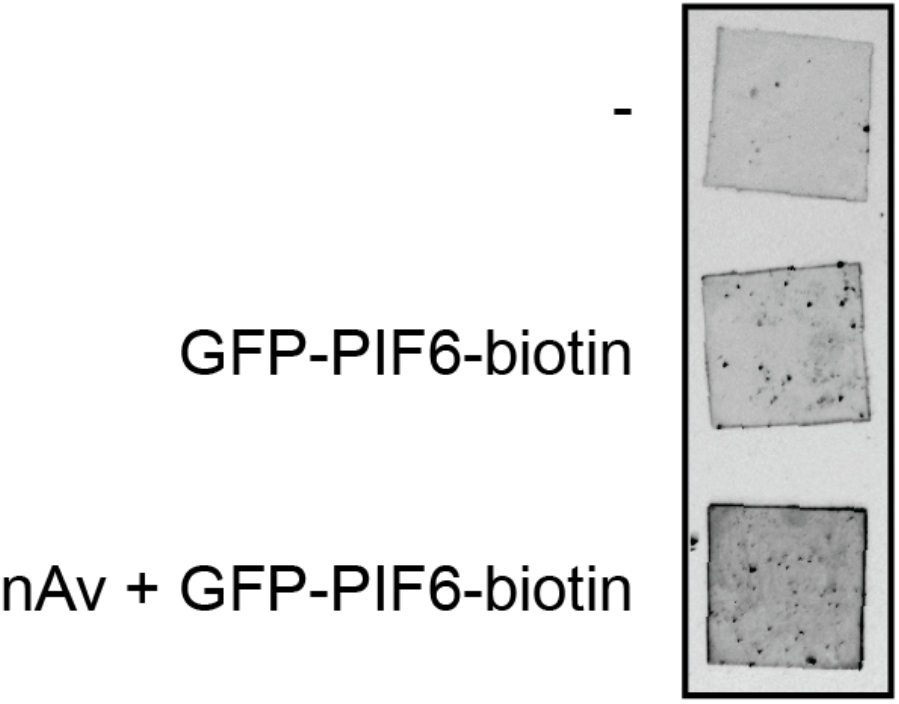
Adsorption Test on PMMA Samples. GFP fluorescence images of PMMA samples. The first sample was only treated with PBS (-) while the second sample was treated with biotinylated GFP-PIF6 only. On the third sample, nAv was first adsorbed followed by blocking with 5% BSA. Then, GFP-PIF6-biotin was added. The images were taken after washing the samples.

**Figure S6.**
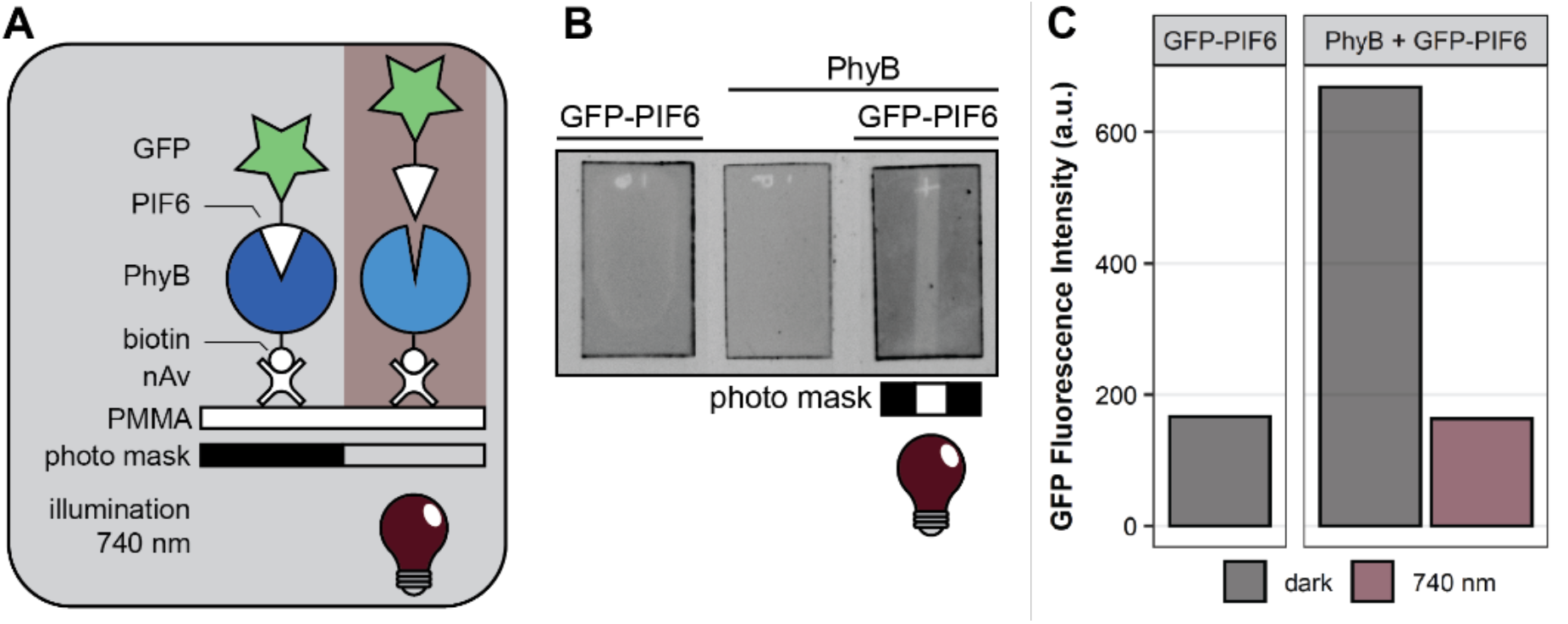
Release Experiments on PMMA Substrate. **A)** Experimental setup of the spatially resolved release of the competitor complex PIF6-GFP from PMMA. Competitor complex was immobilized via biotinylated PhyB on PMMA treated with neutravidin (nAv). The samples were illuminated from below with far-red light (740 nm) through a photomask. **B)** GFP fluorescence images of PMMA samples. The first sample was only treated with GFP-PIF6 and the middle sample was only treated with biotinylated PhyB after nAv treatment and blocking. For the right most sample, first nAv was adsorbed on PMMA, then biotinylated PhyB was added to the. After blocking, GFP-PIF6 was incubated during red light (660 nm) illumination. This sample was then put on a photomask and after addition of buffer to cover the whole area, the sample was illuminated through photomask with far-red light from below for 1 min. The other samples were left in the dark. **C)** Quantification of fluorescence intensity measured from B in a.u. The background fluorescence of the PhyB only sample was subtracted from all samples before plotting. Black bars display the intensity of the not-illuminated samples/areas; the dark-red bar displays the intensity of the illuminated area.

**Figure S7.**
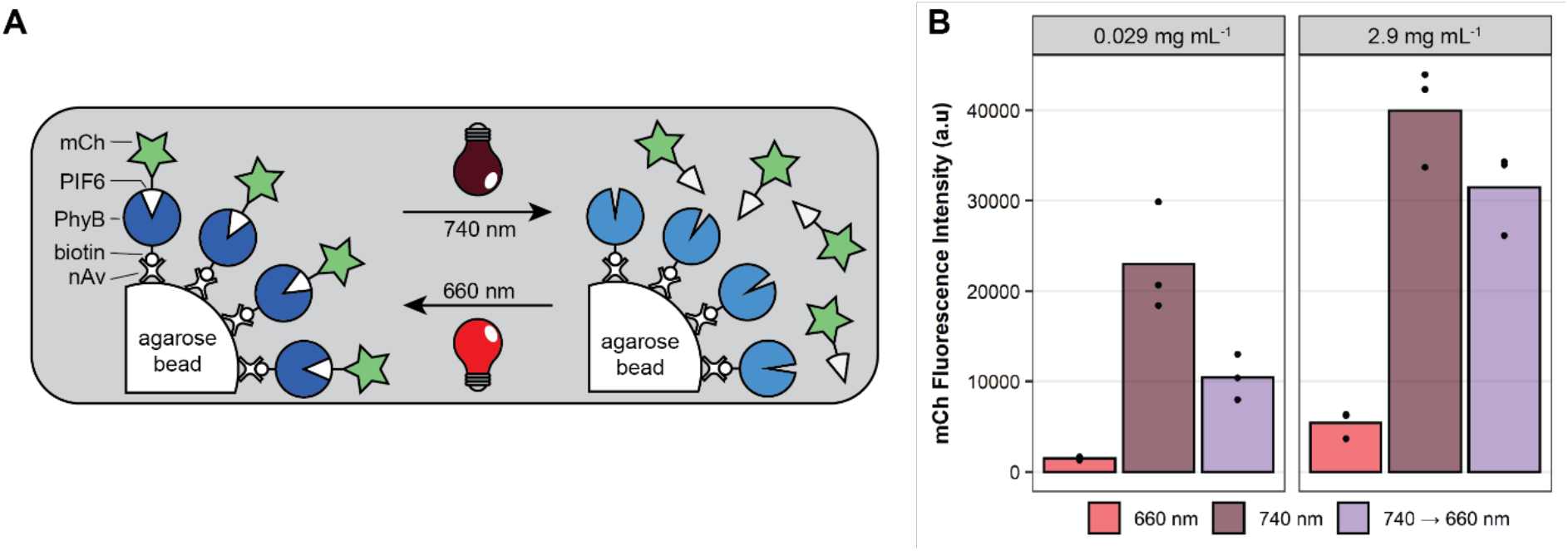
Release and Rebinding Experiments on Beads. **A)** Experimental setup of release and rebinding of the competitor complex PIF6-mCh from and to PhyB-functionalized agarose beads. Illumination with far-red light (740 nm) leads to a release of competitor from PhyB that is coupled to the beads via neutravidin-biotin interaction; during red light (660 nm) illumination mCh-PIF6 and PhyB are associated. **B)** For the preparation, 10 μl PhyB coated beads were incubated with two different concentrations of mCh-PIF6 (0.029 or 2.9 mg ml^−1^) under red light illumination. For this experiment, the loaded beads were incubated in buffer and either illuminated with red or far-red light for 1 h or first with far-red light for 30 min and then with red light for 30 min. The mCh fluorescence intensity was measured in a.u. from the supernatant of the sedimented beads. The background fluorescence of the buffer was subtracted from each value. The means of 3 replicates (dots) are displayed as bars.

## Notes

### Competing Interest Statement

The authors have declared no competing interest.

### Summary of Updates

We have improved the methods section and the language/formatting of our manuscript.

## References

1. Syedmoradi, L. & Gomez, F. A. Paper-based point-of-care testing in disease diagnostics. Bioanalysis 9, 841–843 (2017).

2. Ding, X., Cheung, S. F., Cheng, S. K. L. & Kamei, D. T. Paper-Based Systems for Point-of-Care Biosensing. J. Lab. Autom. 20, 316–333 (2015).

3. Dincer, C., Bruch, R., Kling, A., Dittrich, P. S. & Urban, G. A. Multiplexed Point-of-Care Testing – xPOCT. Trends Biotechnol. 35, 728–742 (2017).

4. Dincer, C. et al. Disposable Sensors in Diagnostics, Food, and Environmental Monitoring. Adv. Mater. 31, 1806739 (2019).

5. Hörner, M. et al. Production of Phytochromes by High-Cell-Density E. coli Fermentation. ACS Synth. Biol. 8, 2442–2450 (2019).

6. Ziegler, T. & Möglich, A. Photoreceptor engineering. Front. Mol. Biosci. 2, 1–25 (2015).

7. Burgie, E. S. & Vierstra, R. D. Phytochromes: An atomic perspective on photoactivation and signaling. Plant Cell 26, 568–4583 (2014).

8. Shimizu-Sato, S., Huq, E., Tepperman, J. M. & Quail, P. H. A light-switchable gene promoter system. Nat. Biotechnol. 20, 1041–1044 (2002).

9. Levskaya, A., Weiner, O. D., Lim, W. A. & Voigt, C. A. Spatiotemporal control of cell signalling using a light-switchable protein interaction. Nature 461, 997–1001 (2009).

10. Beyer, H. M. et al. Generic and reversible opto-trapping of biomolecules. Acta Biomater. 79, 276–282 (2018).

11. Wieland, F. et al. Enhanced Protein Immobilization on Polymers — A Plasma Surface Activation Study. Polymers (Basel). 12, 1–12 (2020).

12. Towbin, H., Staehelin, T. & Gordon, J. Electrophoretic transfer of proteins from polyacrylamide gels to nitrocellulose sheets: procedure and some applications. Proc. Natl. Acad. Sci. 76, 4350–4354 (1979).

13. Hartmeier, W. Immobilized Biocatalysts. (Springer Berlin Heidelberg, 1988). doi:10.1007/978-3-642-73364-2.

14. Chou, J. et al. Porous bead-based diagnostic platforms: Bridging the gaps in healthcare. Sensors (Switzerland) 12, 15467–15499 (2012).

15. Qu, J., Chenier, M., Zhang, Y. & Xu, C. A Microflow Cytometry-Based Agglutination Immunoassay for Point-of-Care Quantitative Detection of SARS-CoV-2 IgM and IgG. Micromachines 12, 433 (2021).

16. Horsley, J. R. et al. Photoswitchable peptide-based ‘on-off’ biosensor for electrochemical detection and control of protein-protein interactions. Biosens. Bioelectron. 118, 188–194 (2018).

17. Adelizzi, B., Gielen, V., Le Saux, T., Dedecker, P. & Jullien, L. Quantitative Model for Reversibly Photoswitchable Sensors. ACS Sensors (2021) doi:10.1021/acssensors.0c02414.

18. Nilsson, B. et al. A synthetic IgG-binding domain based on staphylococcal protein a. Protein Eng. Des. Sel. 1, 107–113 (1987).

19. Guntas, G. et al. Engineering an improved light-induced dimer (iLID) for controlling the localization and activity of signaling proteins. Proc. Natl. Acad. Sci. 112, 112–117 (2015).

20. Ireta-Muñoz, L. A. & Morales-Narváez, E. Smartphone and Paper-Based Fluorescence Reader: A Do It Yourself Approach. Biosensors 10, 60 (2020).

21. Gibson, D. G. et al. Enzymatic assembly of DNA molecules up to several hundred kilobases. Nat. Methods 6, 343–345 (2009).

22. Beyer, H. M. et al. Synthetic Biology Makes Polymer Materials Count. Adv. Mater. 30, 1800472 (2018).

23. Hörner, M., Yousefi, O. S., Schamel, W. & Weber, W. Production, Purification and Characterization of Recombinant Biotinylated Phytochrome B for Extracellular Optogenetics. Bio-Protocol 10, 1–17 (2020).

24. Wieland, F. et al. Enhanced Protein Immobilization on Polymers—A Plasma Surface Activation Study. Polymers (Basel). 12, 104 (2020).

